# Juvenile influenza can impair myelin development and adult behavior through chemokine signaling in mice

**DOI:** 10.64898/2026.08.08.743593

**Authors:** Karen Malacon, Kiarash Shamardani, Sophia Artandi, Lijun Ni, Natalia K. Zernicka-Glover, Abigail Rogers, Belgin Yalçın, Enrique Herrera Castañeda, Theresa Pham, Akiko Iwasaki, Catherine A. Blish, Anna C. Geraghty, Michelle Monje

## Abstract

Brain development, especially developmental myelination, continues through young adulthood. Concordantly, children may be particularly vulnerable to neural-immune challenges. To investigate the consequences of major childhood immune challenges, juvenile mice were exposed to respiratory influenza (H1N1) infection. White matter-specific microglial reactivity accompanied by oligodendrocyte loss was evident until two months following infection. Mice exhibited hyperlocomotion and impaired attention, but not anxiety-like behavior, at one month following infection. Linking the oligodendroglial and behavioral deficits, genetic disruption of oligodendrocyte development at the same juvenile timepoint recapitulated this behavioral phenotype. Microglial reactivity and oligodendrocyte numbers normalized by young adulthood. However, myelin development was disrupted, with persistently decreased myelinated axon density and reduced myelin sheath thickness. Hyperlocomotion resolved, but anxiety-related behaviors emerged at two months after infection. At 6 months, anxiety resolved but cognitive deficits persisted. Elevated CSF chemokines and microglial chemokine expression prompted testing the role of the multi-chemokine receptor CCR3. CCR3 inhibition rescued these cellular and behavioral aberrations after juvenile H1N1 infection. Together, these findings underscore the potential for disruption of myelin development and lasting cognitive and neuropsychiatric sequelae following major immune challenges during the juvenile period and highlight chemokine signaling as an important therapeutic target.

## INTRODUCTION

Systemic inflammation can lead to significant brain dysfunction due to the reactivity of central nervous system (CNS) myeloid cells such as microglia to various immune challenges and the consequent impact on neuron-glial interactions^1–3^ . Respiratory viral infections such as COVID-19 and influenza induce a reactive state in microglia, particularly microglia residing in white matter tracts^2^. This white matter microglial reactivity can impair oligodendroglial and myelin homeostasis^1,2^ and plasticity^4^, as well as the ongoing generation of new neurons in the hippocampus (neurogenesis)^2,5–7^. While the mechanisms by which reactive microglia disrupt oligodendroglial integrity are not fully understood, one contributing factor may be loss of the homeostatic support that microglia provide for oligodendrocytes in states of health and regeneration^8,9^. Reactive microglia can locally secrete cytokines ^6^ and chemokines^2^, as well as induce neurotoxic astrocyte reactivity^10^, contributing to impaired cellular homeostasis and plasticity (for review, see Monje and Iwasaki^11^). Microglia are known to become persistently reactive following exposure to various systemic inflammatory insults, including certain chemotherapy drugs^1,4^, cranial radiation^5,7^, cancer immunotherapies^3^, and low-dose lipopolysaccharide mimicking bacterial infection^6^. This recurrent pattern of microglial reactivity and consequent oligodendroglial dysregulation observed following diverse immune challenges significantly contributes to cognitive impairment, and immune-modulatory strategies targeting microglia or chemokine signaling have been shown to normalize this multicellular dysregulation and rescue cognition in a variety of disease states^1,3,4,6^.

Children are particularly vulnerable to neuroimmune challenges due to ongoing brain development. The first several weeks of life in mice, analogous to the first several years of life in humans, constitute a critical window during which brain architecture is shaped for life^12,13^. During this postnatal period, several key neurodevelopmental processes are active, among which developmental myelination is central and largely occurs after birth. Oligodendrogenesis and myelination are vital for the development, plasticity, and function of neural circuits^14–17^, and accumulating evidence indicates that disruptions in oligodendrocyte function and myelin integrity contribute to neurological and psychiatric disease states^18–20^. For example, experimental disruption of myelin plasticity in early adulthood causes impairment in attention and memory function^4^. In humans, myelin develops throughout the juvenile, adolescent, and young adult periods^21,22^, while in mice, myelin development continues through the first 5 postnatal weeks^14,23,24^. Myelination occurs at predictable times and in specific regions, adhering to well-defined topographical and chronological sequences. Typically, circuits for movement and sensation myelinate first, followed by those supporting higher cognition^25–27^, and large-diameter axons myelinate before smaller axons^28,29^. We hypothesized that inflammatory challenges during this period could disrupt myelin development, leading to long-lasting consequences for cognition and behavior.

While other studies have investigated the impact of prenatal immune challenges, including influenza and the viral mimic poly(I:C)^30–32^, few have explored the long-term effects of juvenile immune insults. Here, we chose to study juvenile exposure to respiratory influenza for several reasons. First, influenza is a common childhood immune challenge that is typically non-neurotropic and has been associated with long-term neurological and neuropsychiatric sequelae (for review, see Smith et al.^33^) Second, we previously showed that a murine model of H1N1 influenza restricted to the respiratory system induces neuroinflammation, including microglial reactivity and elevated CNS cytokines/chemokines and associated oligodendroglial loss in adults^2^. Third, neuroinflammation and the associated oligodendroglial deficits self-resolve in this murine model of influenza by two months after infection^2^, allowing us to test the effects of a discrete epoch of neuroinflammation during the juvenile period on long-term brain function in adulthood.

Considering the ongoing neurodevelopment occurring during the juvenile period, we hypothesized that respiratory influenza infection during early postnatal life may induce neuroinflammation and possibly disruption of myelin development during this crucial developmental stage, which may in turn contribute to putative neurological/neuropsychiatric impairments. Understanding the neural-immune response to such an epoch of neuroinflammation during the juvenile period may suggest avenues for understanding potential later life neurological and neuropsychiatric sequele, and to develop targeted interventions to mitigate long-term neurocognitive effects following major immune challenges in pediatric populations.

## RESULTS

### Respiratory H1N1 Influenza Induces Microglia Reactivity and Disrupts Oligodendrocyte Maturation in the Juvenile Period

To investigate the neuroimmune and cognitive effects of respiratory immune challenges during early postnatal life, we exposed juvenile mice to a relatively mild respiratory influenza A/PR8/34 (H1N1) infection at postnatal day (P)14 via intratracheal administration (**Figure 1A**). Intratracheal inoculation is a superior method for studying the effects of influenza-induced acute lung infection compared with the intranasal route^34^ and it also prevents direct viral exposure of the olfactory bulb^35^. Following infection, juvenile mice showed a transient decrease in expected weight gain between 8 and 10 days post-infection, with weight normalizing by 20 days post-infection (**Figure 1B**). To confirm the previous demonstrations that A/PR8/34 is non-neurotropic^36,37^, we assessed the presence of viral nucleoprotein (NP) in the lung and brain using qRT-PCR (**Figures S1A-S1H**). Virus was detected in the lung at 3 days post-infection and cleared by 2 weeks post-infection (**Figures S1E and S1F**). In contrast, no virus was detected in the brain at either 3 days or 2 weeks post-infection (**Figures S1G and S1H**), confirming that our model of H1N1 infection is non-neurotropic, with infection clearing from the lungs by 2 weeks.

**Figure 1.**
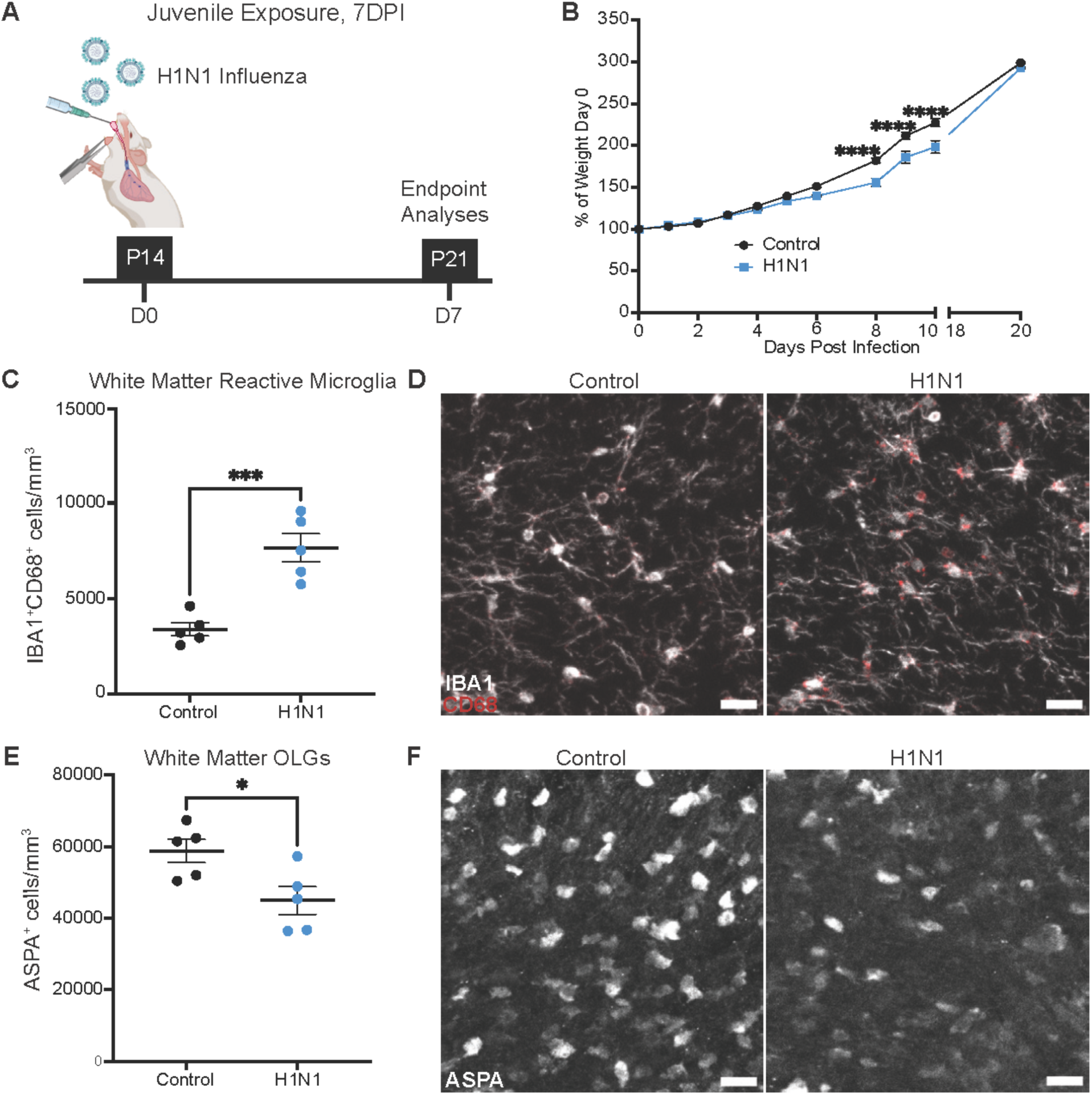
White matter microglial reactivity and oligodendrocyte loss seven days after juvenile H1N1 infection. (A) Schematic of experimental paradigm for mild respiratory H1N1 influenza infection and experimental timeline. Mice were intratracheally inoculated at P14 with PBS or H1N1, and endpoint analyses were performed at P21, 7 days post-infection (DPI). (B) Body weight (% of day 0 weight) of control and H1N1-infected mice. *n* = 10 mice per group. (C) Quantification of white matter reactive microglia (IBA1^+^CD68^+^) 7 days post-infection in the cingulum. *n* = 5 mice per group. (D) Representative confocal micrographs of white matter reactive microglia (IBA1, white; CD68, red) in the cingulum 7 days post-infection. (E) Quantification of white matter mature oligodendrocytes (ASPA^+^) 7 days post-infection in the cingulum. *n* = 5 mice per group. (F) Representative confocal micrographs of white matter mature oligodendrocytes (ASPA, white) in the cingulum 7 days post-infection. Data shown as mean ± SEM; each dot represents an individual mouse. * p < 0.05, *** p < 0.001, analyzed via (C and E) unpaired two-tailed t-test or (B) two-way ANOVA with multiple comparisons. (D and F) Scale bars, 40 μm.

Various CNS and systemic insults can cause reactivity of microglia, the resident immune cells of the CNS, particularly in white matter regions^1–3^. We evaluated microglial numbers and reactivity 7 days following H1N1 infection (P21), a timepoint during active infection, using the pan-microglial marker Iba1 and the lysosomal marker CD68 in the cortex, subcortical white matter, and hippocampal dentate gyrus white matter. Total Iba1+ microglial numbers were unchanged in all regions assessed (**Figures S2A-S2C**). Microglial reactivity, defined as Iba1 and CD68 co-expression, was increased in subcortical and hippocampal white matter 7 days after infection, but not in the cortical grey matter (**Figures 1C, 1D, S2D, and S2E**).

Microglial reactivity is known to disrupt oligodendroglial homeostasis and plasticity^1,4^. To assess oligodendroglial integrity following H1N1 infection, we quantified oligodendrocyte precursor cells (OPCs) and mature oligodendrocytes in the cortex and subcortical white matter. While no changes in OPC numbers 7 days post-infection were found in either region (**Figures S2F and S2G**), a decrease in mature oligodendrocytes was evident in the subcortical white matter (**Figures 1E and 1F**). Concordant with the white matter-selective pattern of microglial reactivity, mature oligodendrocyte number was not altered in the cortex 7 days post-infection (**Figure S2H**).

### Impaired Cognition Amidst Persistent Glial Dysregulation Four Weeks Post-Infection

Given that microglial reactivity and mature oligodendrocyte loss in white matter are associated with cognitive deficits^3,4^, and considering evidence of long-lasting attention and memory impairments following systemic insults^38–40^ (for reviews, see Panagea et al.^41^ and Calsavara et al.^42^), we conducted a series of behavioral tests in the young adult period (P42), 4 weeks post-infection, when H1N1 infection had cleared from the lungs (**Figures 2A and S1A-H**). To assess anxiety-like behavior and locomotion, we utilized the open field test (**Figures 2B and S3A**). No differences between previously H1N1-infected and control groups were found in time spent in the center of the open field or in the total number of fecal boli— both measures of anxiety-like behavior in mice—at this young adult timepoint (**Figures 2D and S3B**). However, mice previously infected with H1N1 exhibited increased total distance traveled (**Figure 2E**), demonstrating heightened locomotion. The Novel Object Recognition Task (NORT) was used with a short-interval (5-minutes) to weight the test towards testing attention^43,44^ (**Figure 2C**). Young adult mice with previous H1N1 infection in the juvenile period exhibited impaired recognition of the novel object compared with the familiar object in NORT, indicating attention and/or short-term memory deficits at 4 weeks post-infection (**Figure 2F**).

**Figure 2.**
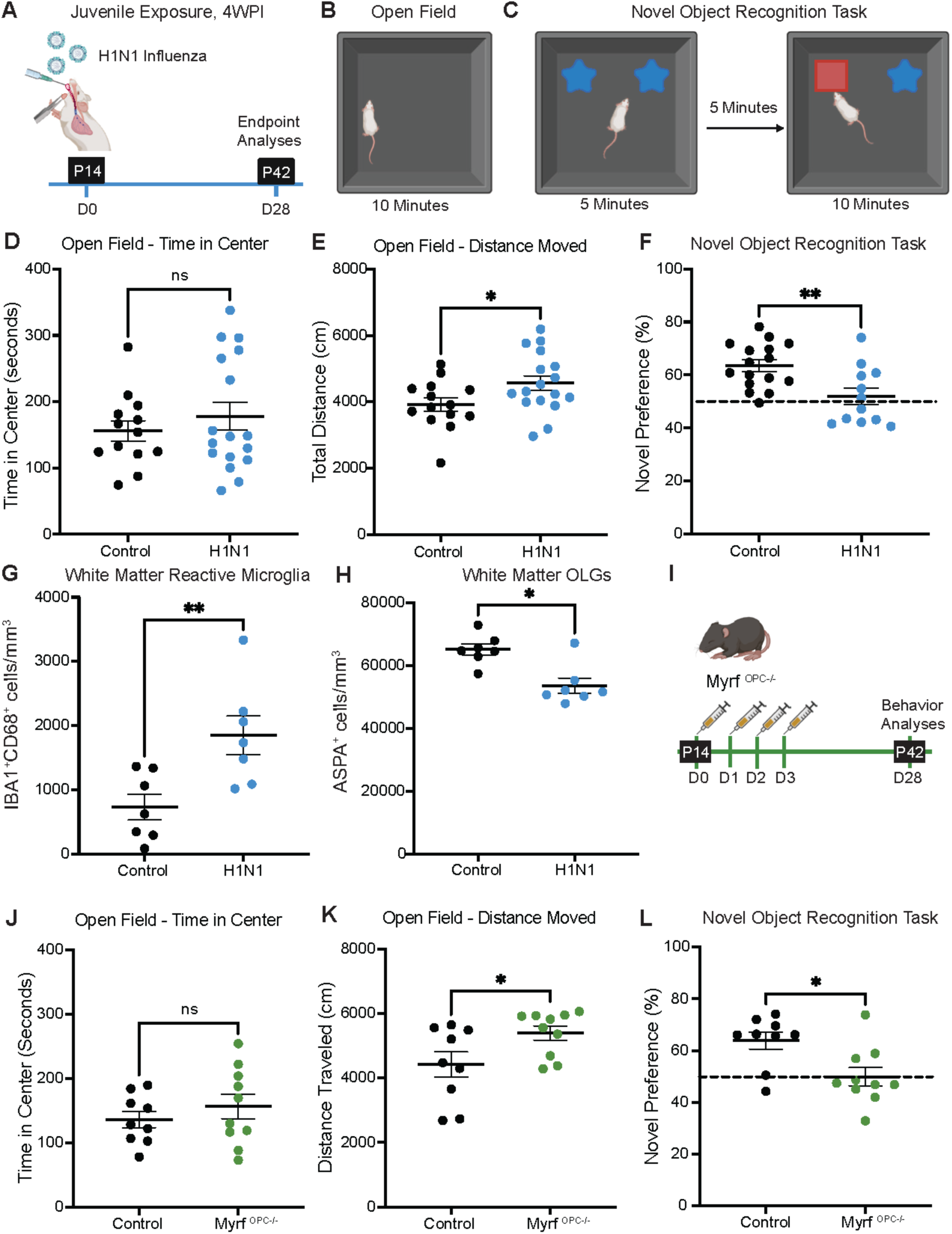
Persistent glial dysregulation accompanies hyperlocomotion and cognitive deficits four weeks post juvenile H1N1 infection. (A) Schematic illustration of experimental timeline. Mice were intratracheally inoculated at P14 with PBS or H1N1, and endpoint analyses were performed at P42, 4 weeks post-infection (WPI). (B) Schematic illustration of the open field to assess anxiety-like behavior and locomotion. (C) Schematic illustration of the novel object recognition task (NORT) of attention and short-term memory. Mice are introduced to two identical objects and tested 5 minutes after for attention by switching one of the objects with a novel object. Novel preference is defined as the percentage of time spent interacting with the novel object relative to total time spent interacting with either object. (D) Time spent in center of the open field over 10-minute testing period. Open field performed 4 weeks post-infection. *n* = 13 control, *n* = 17 H1N1 mice. (E) Total distance moved in the open field over 10-minute testing period. Open field performed 4 weeks post-infection. *n* = 14 control, *n* = 17 H1N1 mice. (F) Novel preference for NORT performed 4 weeks post-infection. *n* = 16 control, *n* = 12 H1N1 mice. (G) Quantification of white matter reactive microglia (IBA1^+^CD68^+^) 4 weeks post-infection in the cingulum. *n* = 7 mice per group. (H) Quantification of white matter mature oligodendrocytes (ASPA^+^) 4 weeks post-infection in the cingulum. *n* = 7 mice per group. (I) Schematic illustration of experimental timeline. *Myrf^OPC-/-^* and *Myrf*-wild-type control were injected with tamoxifen (i.p.) for four consecutive days starting at P14. Behavioral analyses were performed at P42, 4 weeks post-injection. (J) Total distance moved in the open field over 10-minute testing period. Open field performed at P42, 4 weeks post-injection. *n* = 7 control, *n* = 6 *Myrf^OPC-/-^* mice. (K) Novel preference for NORT performed at P42, 4 weeks post-injection. *n* = 7 control, *n* = 6 *Myrf^OPC-/-^* mice. (L) Time spent in center of the open field over 10-minute testing period. Open field performed at P42, 4 weeks post-injection. *n* = 7 control, *n* = 6 *Myrf^OPC-/-^* mice. Data shown as mean ± SEM; each dot represents an individual mouse. ns: p > 0.05, * p < 0.05, ** p < 0.01, analyzed via (E-G, and J) unpaired two-tailed t-test or (D, H, K, and L) Mann-Whitney U test for datasets that failed the Shapiro-Wilk test.

To further explore cognitive performance, particularly hippocampal-related spatial working memory, we employed the spontaneous alternation T-maze test (**Figure S3C**), a well-established measure of spatial memory requiring intact hippocampal function^45,46^. H1N1-infected mice were unable to perform adequately in the T-maze test at 4 weeks post-infection, suggesting compromised spatial memory (**Figure S3D**). To explore possible deficits in social behavior, we used the three-chamber sociability test to assess preference for social versus non-social stimuli (**Figure S3E**). No differences in social preference were observed between the groups, indicating preserved social behavior after juvenile H1N1 respiratory infection (**Figure S3F**).

We then examined whether the glial dysregulation observed at 7 days following infection persisted at 4 weeks post-H1N1 infection. At this timepoint, the total number of microglia remained non-elevated in all regions examined, with a mild reduction in microglial numbers in the subcortical white matter and cortex in previously H1N1-infected mice (**Figures S4A-S4C**). Increased microglial reactivity was still evident in the subcortical and hippocampal white matter of previously H1N1-infected mice (**Figures 2G, S4D, and S4E**). Microglial reactivity in the cortex remained unaltered and similar to control mice (**Figure S4F**). Mature oligodendrocytes remained decreased in the subcortical white matter at this 4-week timepoint following H1N1 influenza infection, with no changes in OPC numbers (**Figures 2H, S4G, and S4H**), indicating ongoing disruption of the white matter oligodendrocyte population. OPCs and mature oligodendrocytes in the cortex remained unaffected by H1N1 infection (**Figures S4I and S4J**).

Subcortical white matter oligodendrocyte homeostasis and plasticity are implicated in attention and memory function^3,4,4,15^. We hypothesized that disruption of white matter oligodendroglial development underlie the impairment in attention and elevated motor activity observed in young adult mice following juvenile H1N1 influenza. To test this hypothesis, we used a conditional genetic mouse model to disrupt oligodrodroglial development beginning at the same postnatal timepoint, P14. Myelin regulatory factor (*Myrf*), a transcription factor required for full oligodendrocyte differentiation, was conditionally deleted from oligodendroglial lineage precursor cells at P14 in mice expressing floxed *Myrf* and tamoxifen-inducible Cre under the OPC-selective promoter *Pdgfra* (*Pdgfra*::Cre-ER; Myrf^lox/lox^). Following deletion of *Myrf* in OPCs with tamoxifen administration, OPCs cannot differentiate into new oligodendrocytes and undergo apoptosis, while pre-existing oligodendrocytes and myelin are unaffected^47^. At 4-weeks following juvenile (P14) disruption of oligodendrocyte generation, mice exhibited impaired performance in the novel object recognition test and hyperlocomotion, with unaffected time spent in the center of the open field and no deficits in social interactions nor T-maze (**Figures 2J-2L, S3G, and S3I**). Taken together, these results support the interpretation that disruption of oligodendroglial development results in the attentional deficits and elevated motor activity observed in young adulthood after juvenile influenza exposure.

### Long-term Behavioral and Myelin Deficits 8 Weeks Following H1N1 Infection

To determine whether the behavioral deficits and glial dysregulation observed at 4 weeks post-infection persisted long-term following juvenile H1N1 infection, we conducted the same behavioral assays and histological analyses at 8 weeks post-infection (P70), when mice were adults (**Figure 3A**). In contrast to the 4-week timepoint, previously H1N1-infected mice spent less time in the center in the open field test and exhibited an increased number of fecal boli (**Figure 3B, S5A, and S5B**), both measures indicating anxiety-like behavior in the adult period. No differences in the total distance traveled were evident between control and previously H1N1-infected mice, suggesting that the aberrantly elevated locomotion observed at the 4-week, young adult timepoint resolved by this stage (**Figure 3C**). Cognitive deficits persisted at 8 weeks post-infection, as mice infected with H1N1 in the juvenile period continued to perform poorly in the NORT, indicating ongoing attention and/or short-term memory challenges (**Figure 3D**). Similarly, deficits in the T-maze test persisted, reflecting continued impairment in spatial working memory (**Figure S5C**). However, sociability remained intact, with no observed deficits in social behavior (**Figure S5D**).

**Figure 3.**
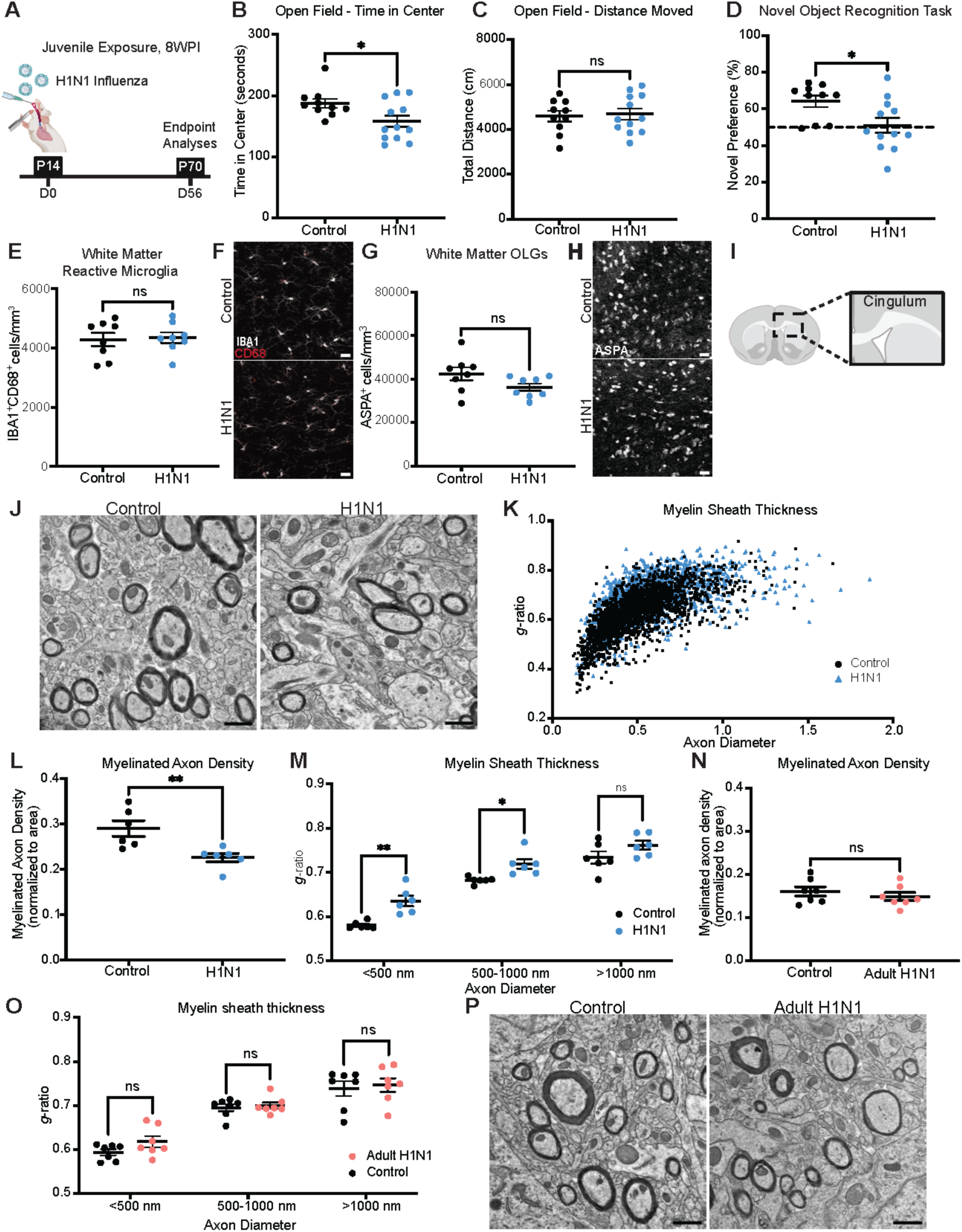
Myelin deficits and sustained cognitive impairments eight weeks following juvenile H1N1 exposure. (A) Schematic illustration of experimental timeline. Mice were intratracheally inoculated at P14 with PBS or H1N1, and endpoint analyses were performed at P70, 8 weeks post-infection (WPI). (B) Time spent in center of the open field over 10-minute testing period. Open field performed 8 weeks post-infection. *n* = 10 control, *n* = 12 H1N1 mice. (C) Total distance moved in the open field over 10-minute testing period. Open field performed 8 weeks post-infection. *n* = 10 control, *n* = 12 H1N1 mice. (D) Novel preference for NORT performed 8 weeks post-infection. *n* = 10 control, *n* = 12 H1N1 mice. (E) Quantification of white matter reactive microglia (IBA1^+^CD68^+^) 8 weeks post-infection in the cingulum. *n* = 8 mice per group. (F) Representative confocal micrographs of white matter reactive microglia (IBA1, white; CD68, red) in the cingulum 8 weeks post-infection. (G) Quantification of white matter mature oligodendrocytes (ASPA^+^) 8 weeks post-infection in the cingulum. *n* = 8 mice per group. (H) Representative confocal micrographs of white matter mature oligodendrocytes (ASPA, white) in the cingulum 8 weeks post-infection. (I) Schematic depicting the cingulum region of the corpus callosum where transmission electron microcopy (TEM) was performed 8 weeks post-infection in mice infected during the juvenile period (J-M) and mice infected during adulthood (N-P). (J) Representative TEM images of mice infected with H1N1 during the juvenile period 8 weeks post-infection at the level of the cingulum of the corpus callosum in cross-section. Myelinated axons visible as electron-dense sheaths encircling axons. (K) Scatter plots of *g*-ratio as a function of axon diameter 8 weeks post-infection for control axons (black dots) or H1N1 axons (blue triangles). A single point indicates the *g*-ratio for a single axon. Approximately 250-400 axons were quantified for each animal. *n* = 6 mice per group. (L) Quantification of myelinated axon density. *n* = 6 mice per group. (M) *G*-ratio relative to small (<500 nm), medium (500-1000 nm), and large (>1000 nm) caliber axons for control (black points) or H1N1 (blue points) treated mice. *n* = 6 mice per group. (N) Quantification of myelinated axon density. *n* = 7 mice per group. (O) *G*-ratio relative to small (<500 nm), medium (500-1000 nm), and large (>1000 nm) caliber axons for control (black points) or H1N1 (orange points) treated mice. *n* = 7 mice per group. (P) Representative TEM images of mice infected with H1N1 during adulthood 8 weeks post-infection at the level of the cingulum of the corpus callosum in cross-section. Myelinated axons visible as electron-dense sheaths encircling axons. Data shown as mean ± SEM (B-E, G, L, and N); each dot represents an individual mouse. ns: p > 0.05, * p < 0.05, ** p < 0.01, analyzed via (B, C, E, G, L, and N) unpaired two-tailed t-test, (D) Mann-Whitney U test for datasets that failed the Shapiro-Wilk test, or (M and O) unpaired two-tailed t-test with correction with multiple comparisons. Scale bars, (F and H) 40 μm, (J and P) 1 μm.

Microglial reactivity in the subcortical white matter had normalized by 8 weeks following juvenile H1N1 influenza infection **(Figures 3E and 3F)**, concordant with the resolution of microglial reactivity two months after adult H1N1 influenza infection^2^. No differences in total microglial numbers between groups were found in any brain region assessed at this 8-week time point (**Figures S5E-S5G**). However, microglial reactivity persisted in the hippocampal white matter at 8 weeks following juvenile H1N1 influenza infection (**Figure S5H**), also concordant with findings after adult H1N1 infection^2^. In the cortex, microglial reactivity remained unaltered (**Figure S5I**). At 8 weeks following juvenile H1N1 infection, the number of mature oligodendrocytes in subcortical white matter normalized to the levels observed in control mice, with OPC numbers remaining unaltered between groups (**Figures 3G, 3H, and S5J**). The number of OPCs and mature oligodendrocytes in the cortex remained unaltered (**Figures S5K and S5L**).

Despite the normalization of mature oligodendrocyte numbers by 8 weeks following juvenile H1N1 infection, the persistent cognitive deficits observed suggest potential disruption of myelin development. As developmental myelination chiefly occurs in the postnatal period, we suspected that myelin development could have been disrupted by this juvenile immune challenge. To investigate changes in myelin ultrastructure, we performed transmission electron microscopy (TEM) of the subcortical white matter (cingulum of the corpus callosum; **Figure 3I**). This revealed a 28% decrease in myelinated axon density in the subcortical white matter at 8 weeks following juvenile H1N1 infection (**Figires 3J-3L and S6A**). This reduction was particularly evident in small diameter (<500 nm) myelinated axons (**Figure S6B**), which are known to myelinate later in life compared to larger diameter axons that myelinate earlier ^28,29^. Furthermore, we observed an overall decrease in myelin sheath thickness (increased *g*-ratio), with thinner myelin sheaths specifically associated with small and medium-sized axons (500-1000 nm; **Figures 3K, 3M, and S6C**). In contrast, H1N1 influenza respiratory infection in adult mice did not result in altered myelination 8 weeks later (**Figures 3N-3P and S6D-S6F**). These findings suggest that juvenile respiratory immune challenge can disrupt myelin development, potentially contributing to the observed alterations in adult behavior.

### Persistent Cognitive and Myelin Impairments 6 Months Following Infection

Taking into account the myelin ultrastructure changes observed at 8 weeks post-infection and reports of prolonged cognitive impairment in children following respiratory immune challenges ^48–50^, we extended our study to follow a cohort of mice inoculated with H1N1 or vehicle at P14 for 6 months following infection (**Figures 4A and S7A**). At P194, we found no differences between H1N1-infected and control mice in time spent in the center of the open field, number of fecal boli, nor total distance traveled in the open field test (**Figures 4B, 4C, and S7B**). Despite the normalization of anxiety-like behaviors and hyperlocomotion, mice infected with H1N1 in the juvenile period continued to exhibit cognitive deficits in the NORT test of attention and short-term memory (**Figure 4D**). Concordantly, the deficits in subcortical myelination persisted at 6 months (**Figures 4E-4G, S6E, and S6F**). However, mice no longer exhibited deficits in the spontaneous alternation T-maze test (**Figure S7C**), suggesting recovery of spatial working memory over time. Additionally, we continued to find preserved social behavior, with no differences in social preference in the three-chamber sociability test (**Figure S7D**).

**Figure 4.**
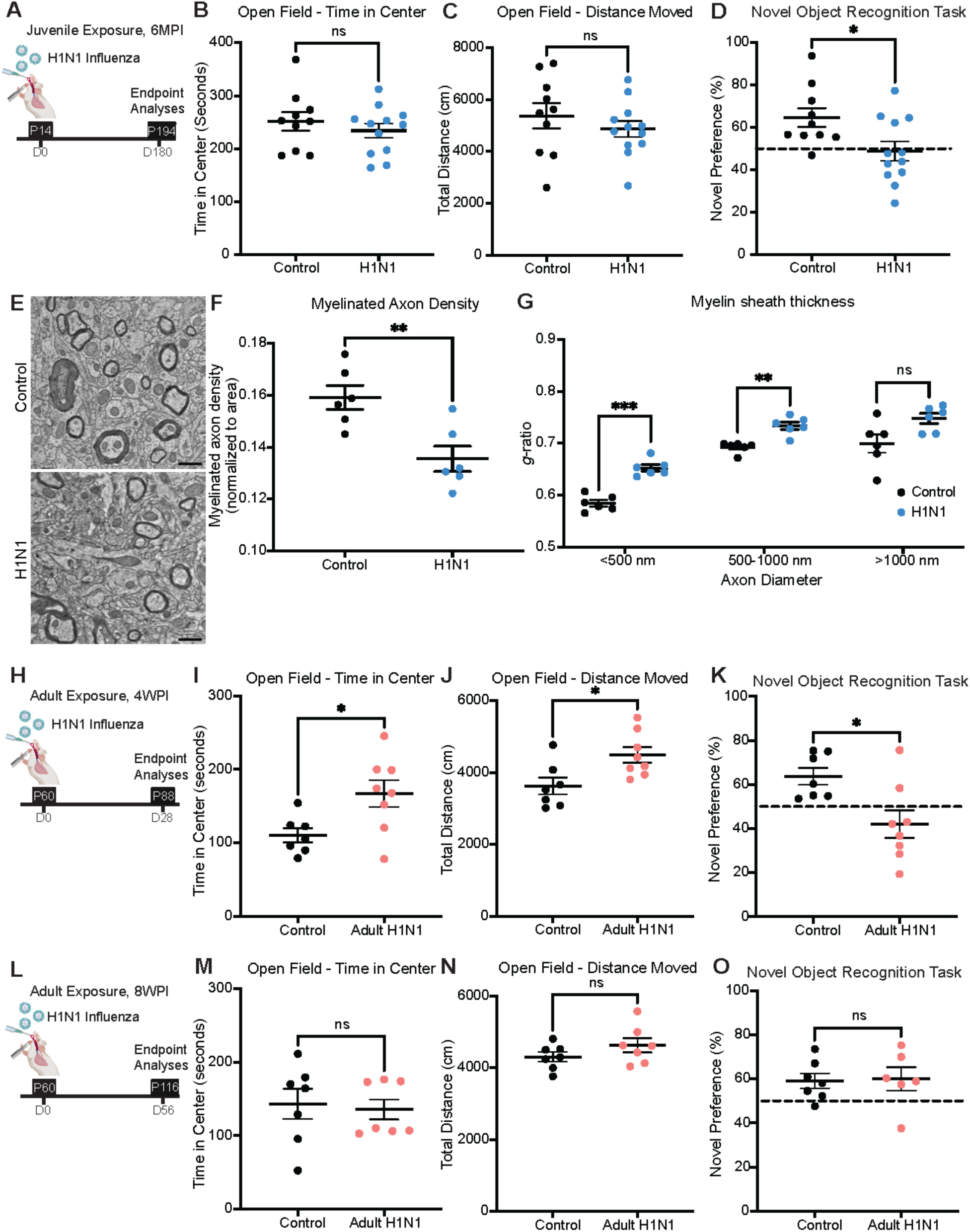
Cognitive deficits resolve in adult-infected mice but cognitive and myelin deficits persist long-term after juvenile H1N1 infection. (A) Schematic illustration of experimental timeline. Mice were intratracheally inoculated at P14 with PBS or H1N1, and endpoint analyses were performed at P194, 6 months post-infection (MPI). (B) Time spent in center of the open field over 10-minute testing period. Open field performed 6 months post-infection. *n* = 10 control, *n* = 12 H1N1 mice. (C) Total distance moved in the open field over 10-minute testing period. Open field performed 6 months post-infection. *n* = 10 control, *n* = 12 H1N1 mice. (D) Novel preference for NORT performed 6 months post-infection. *n* = 10 control, *n* = 12 H1N1 mice. (E) Representative TEM images of mice infected with H1N1 during the juvenile period 6 months post-infection at the level of the cingulum of the corpus callosum in cross-section. Myelinated axons visible as electron-dense sheaths encircling axons. (F) Quantification of myelinated axon density. *n* = 6 mice per group. (G) *G*-ratio relative to small (<500 nm), medium (500-1000 nm), and large (>1000 nm) caliber axons for control (black points) or H1N1 (blue points) treated mice. *n* = 6 mice per group. (H) Schematic illustration of experimental timeline. Mice were intratracheally inoculated at P60 with PBS or H1N1, and endpoint analyses were performed at P88, 4 weeks post-infection. (I) Time spent in center of the open field over 10-minute testing period. Open field performed 4 weeks post-infection. *n* = 7 control, *n* = 8 H1N1 mice. (J) Total distance moved in the open field over 10-minute testing period. Open field performed 4 weeks post-infection. *n* = 7 control, *n* = 8 H1N1 mice. (K) Novel preference for NORT performed 4 weeks post-infection. *n* = 7 mice per group. (L) Schematic illustration of experimental timeline. Mice were intratracheally inoculated at P60 with PBS or H1N1, and endpoint analyses were performed at P116, 8 weeks post-infection. (M) Time spent in center of the open field over 10-minute testing period. Open field performed 8 weeks post-infection. *n* = 7 mice per group. (N) Total distance moved in the open field over 10-minute testing period. Open field performed 8 weeks post-infection. n = 7 mice per group. (O) Novel preference for NORT performed 8 weeks post-infection. *n* = 7 control, *n* = 6 H1N1 mice. Data shown as mean ± SEM; each dot represents an individual mouse. ns: p > 0.05, * p < 0.05, analyzed via (B-D, F, I-K, N, and O) unpaired two-tailed t-test, (G) unpaired-t tests with correction for multiple comparisons, or (M) Mann-Whitney U test for datasets that failed the Shapiro-Wilk test. Scale bar (E), 1 μm.

### Cognitive Deficits at 4 Weeks Post-Infection in Adult H1N1 Infected Mice Normalizes by 8 Weeks Post-Infection

We wanted to determine whether the cognitive deficits observed following juvenile H1N1 infection at P14 were specific to this developmental stage or if similar impairments could occur following infection during adulthood. Previous work has shown that adult mice infected with respiratory H1N1 exhibit glial dysregulation similar to that observed in our P14 model, with microglial reactivity in subcortical and hippocampal white matter and a loss of mature oligodendrocytes in the subcortical white matter at 7 days post-infection. These glial populations normalize by seven weeks post-infection^2^. To investigate the behavioral consequences of adult H1N1 influenza infection, we intratracheally infected two-month-old (P60) CD1 mice with a mild dose of H1N1 and conducted behavioral testing 4 weeks post-infection (P88; **Figures 4H, S7G, and S7H**). Adult mice previously infected with H1N1 influenza displayed increased time spent in the center of the open field, no change in the number of fecal boli, and increased distance traveled (**Figures 4I, 4J, and S7I**). This hyperlocomotion phenotype mirrors what we observed in mice infected with influenza in the juvenile period at 4 weeks post-infection. The increased time spent in the center was interpreted as a result of heightened locomotion rather than reduced anxiety-like behavior, given the unchanged number of fecal boli in the open field compared to control mice. Mice infected during adulthood also exhibited cognitive impairment in the NORT (**Figure 4K**), suggesting attention and memory challenges similar to those seen with juvenile infection. However, unlike mice infected at P14, adult mice showed no deficits in the spontaneous alternation T-maze test (**Figure S7J**). No changes in social behavior were observed at this timepoint in mice infected during adulthood (**Figure S7K**). To test whether the observed behavioral effects in cognition and hyperlocomotion persisted long-term, we repeated these behavioral tests at 8 weeks post-infection (P116; **Figures 4L and S7L**). At 8 weeks following adult H1N1 influenza infection, we found no differences in time spent in the center, number of fecal boli, or distance traveled in the open field (**Figures 4M, 4N, and S7M)**, indicating normalization of hyperlocomotion. Cognitive impairment in the NORT was no longer evident (**Figure 4O**), and mice continued to perform well in the T-maze and sociability tests (**Fgures S7N and S7O**). These findings suggest that while adult mice infected with H1N1 exhibit mild behavioral disruptions at 4 weeks post-infection, these deficits resolve by 8 weeks post-infection – concordant with the normal myelination evident in mice 8 weeks following adult infection described above. In contrast, juvenile mice infected at P14 exhibit persistent behavioral disruptions, highlighting the heightened vulnerability of the developing brain to immune challenges such as H1N1 influenza.

### Single-Nucleus Transcriptomic Profiling After Juvenile H1N1 Infection

To characterize transcriptomic changes in microglia and oligodendroglia following juvenile H1N1 exposure, given previous identification of disease-associated microglial states across neurodegenerative and inflammatory conditions^51,52^, we performed single-nucleus RNA sequencing of white matter in the cingulum at 7 days after respiratory H1N1 infection (26,653 single nuclei from control mice and 27,573 single nuclei from H1N1-infected mice; **Figures 5A and S8A**). We identified the expected cell types, including various glial and neuronal classes (**Figure S8B**). Within the microglia cluster, gene ontology (GO) enrichment analysis revealed significant enrichment of several biological pathways associated with strong immune responses in H1N1-infected mice. Specifically, microglia from H1N1-infected mice displayed upregulation of pathways related to the innate immune response and antigen processing, along with downregulation of pathways involved in regulation of synapse structural plasticity (**Figure S8C**). These changes are consistent with a shift toward an antigen-presenting, pro-inflammatory state that may be less supportive of normal synaptic remodeling. In line with this, microglia from H1N1-infected mice exhibited increased expression of major histocompatibility complex class I (MHC I) implicated in antigen presentation, including *H2-K1* and *H2-Q4* (**Figure 5B**). We next investigated whether microglia exhibited previously defined inflammatory and immune-related gene-state signatures following H1N1 infection^2^. Microglia from H1N1-infected mice had higher scores for chemokine genes (e.g. *Ccl2, Ccl4, Ccl3*, and *Cxcl10*; **Figure 5C**), inflammation-related genes (e.g. *Tlr2, Il1a,* and *Tnfaip2*; **Figure 5D**), and interferon-responsive genes (e.g. *Ifit3, Ifit2,* and *Isg15*; **Figure 5E**), consistent with ongoing antiviral responses and an inflammatory-reactive response.

**Figure 5.**
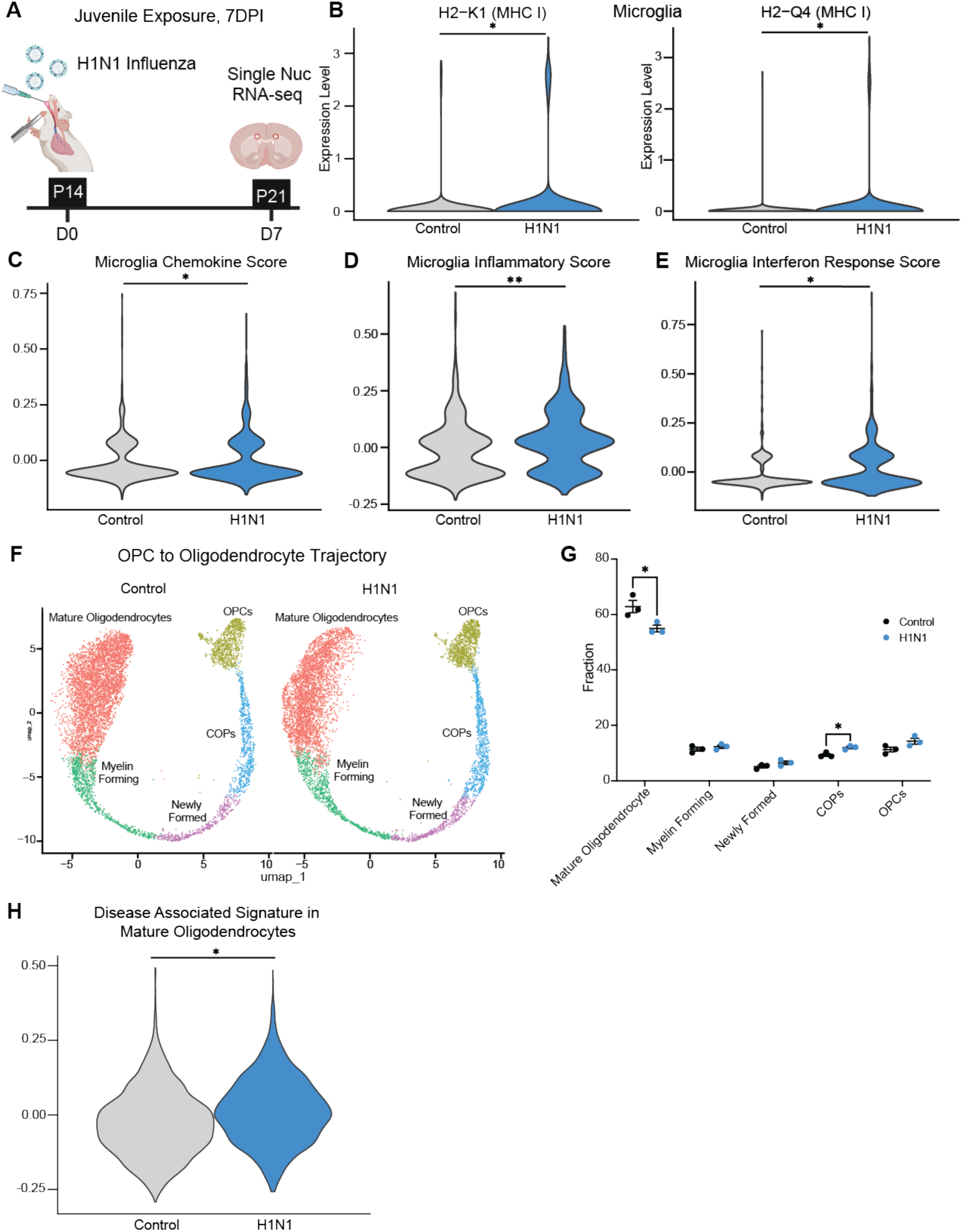
White matter enriched microglial and oligodendroglial single-nuclei transcriptomic profiles 7 days after juvenile exposure to H1N1. (A) Schematic illustration of experimental timeline. Mice were intratracheally inoculated at P14 with PBS or H1N1, and single-nucleus RNA sequencing of the cingulum was performed at P21, 7 days post-infection (DPI). (B) Violin plots illustrating expression of major histocompatibility complex class I (MHC I) genes, H2-K1 (left) and H2-Q4 (right), upregulated in microglia from H1N1 samples compared with controls. (C) Violin plot illustrating chemokine gene expression scores in microglia from control and H1N1 samples. (D) Violin plot illustrating inflammatory gene expression scores in microglia from control and H1N1 samples. (E) Violin plot illustrating interferon response gene expression scores in microglia from control and H1N1 samples. (F) UMAP plot of oligodendroglia trajectory from OPCs to mature oligodendrocytes, colored by cluster, split by control (left, *n* = 3 mice per group, 7,189 nuclei) and H1N1 (right, *n* = 3 mice per group, 6,808 nuclei) samples. (G) Percentage of stages of differentiation of oligodendroglia in the cingulum. *n* = 3 mice per group. (H) Violin plot illustrating expression of disease-associated oligodendrocyte (DOL) gene signature in mature oligodendrocytes from control and H1N1 samples. For statistical analysis in B-E and H, each sample was pseudobulked. * p < 0.05, ** p < 0.01, analyzed via (B-E and H) unpaired two-tailed t-test or (G) one-way ANOVA.

To evaluate whether H1N1 infection and the consequent immune response altered oligodendrocyte lineage progression, we identified established subpopulations of oligodendroglia^53^ (**Figure 5F**). H1N1 infection shifted oligodendroglial differentiation states, with fewer mature oligodendrocytes and relatively more PDGFRα^-^ differentiation-committed oligodendrocyte precursor cells (COPs) compared to controls (**Figure 5G**), suggesting delayed differentiation or potential accumulation of less mature cells in the lineage. GO enrichment analysis within the OPC cluster revealed suppression of DNA replication and cell-cycle division pathways in OPCs from H1N1-infected mice versus controls (**Figure S8D**), indicating reduced proliferative capacity. Such changes could impede oligodendrocyte maturation and myelination. We next compared the mature oligodendrocyte cluster to a previously described disease-associated oligodendrocyte (DOL) signature shared across several CNS pathologies, including Alzheimer’s disease and experimental autoimmune encephalomyelitis^54^. Mature oligodendrocytes from H1N1-infected mice exhibited an upregulated disease-associated oligodendrocyte signature compared to healthy controls (**Figure 5H**).

### Chemokine Receptor Inhibition Rescues Oligodendroglial Dysregulation and Behavioral Aberrations

To further explore cytokine and chemokine profiles following juvenile H1N1 infection, we collected cerebrospinal fluid (CSF) 4 weeks post-infection and measured the levels of various cytokines and chemokines. At this time point, we observed that CSF cytokine and chemokine levels were elevated in previously H1N1-infected mice compared to control mice (**Figure 6A**). Notably, IL-22, IL-25/IL-17, CCL5, and CCL11 CSF levels were significantly elevated at 4-weeks following H1N1 infection. Given the increase in CSF chemokine levels and the increased chemokine expression score in microglia from H1N1-infected mice, we explored whether targeting a chemokine receptor could mitigate the neurobiological effects of H1N1 infection. We focused on CCR3, a key receptor for CCL5 and CCL11, elevated in the CSF of previously H1N1-infected mice and in light of our previous discovery that targeting CCR3 mitigates the neurobiological effects of another immune challenge, CAR T cell immunotherapy^3^. To test the hypothesis that CCR3 chemokine signaling may be central to the respiratory immune challenge-induced cellular and behavioral dysregulation observed, we administered a blood-brain-barrier (BBB)-permeable CCR3 inhibitor (SB-328437) from day 7 to day 28 following juvenile H1N1 infection at P14 (**Figure 6B**). Behavioral testing at 4 weeks post-infection revealed that CCR3 inhibition normalized hyperlocomotion in the open field test (**Figure 6C**). This treatment did not affect the total time spent in the center of the open field (**fig. S8A**). Cognitive performance in the NORT was also rescued in CCR3 inhibitor-treated animals following juvenile H1N1 infection (**Figure 6D**). However, CCR3 inhibition did not rescue deficits in the T-maze test; social behavior remained unaffected (**Figures S8B and S8C**).

**Figure 6.**
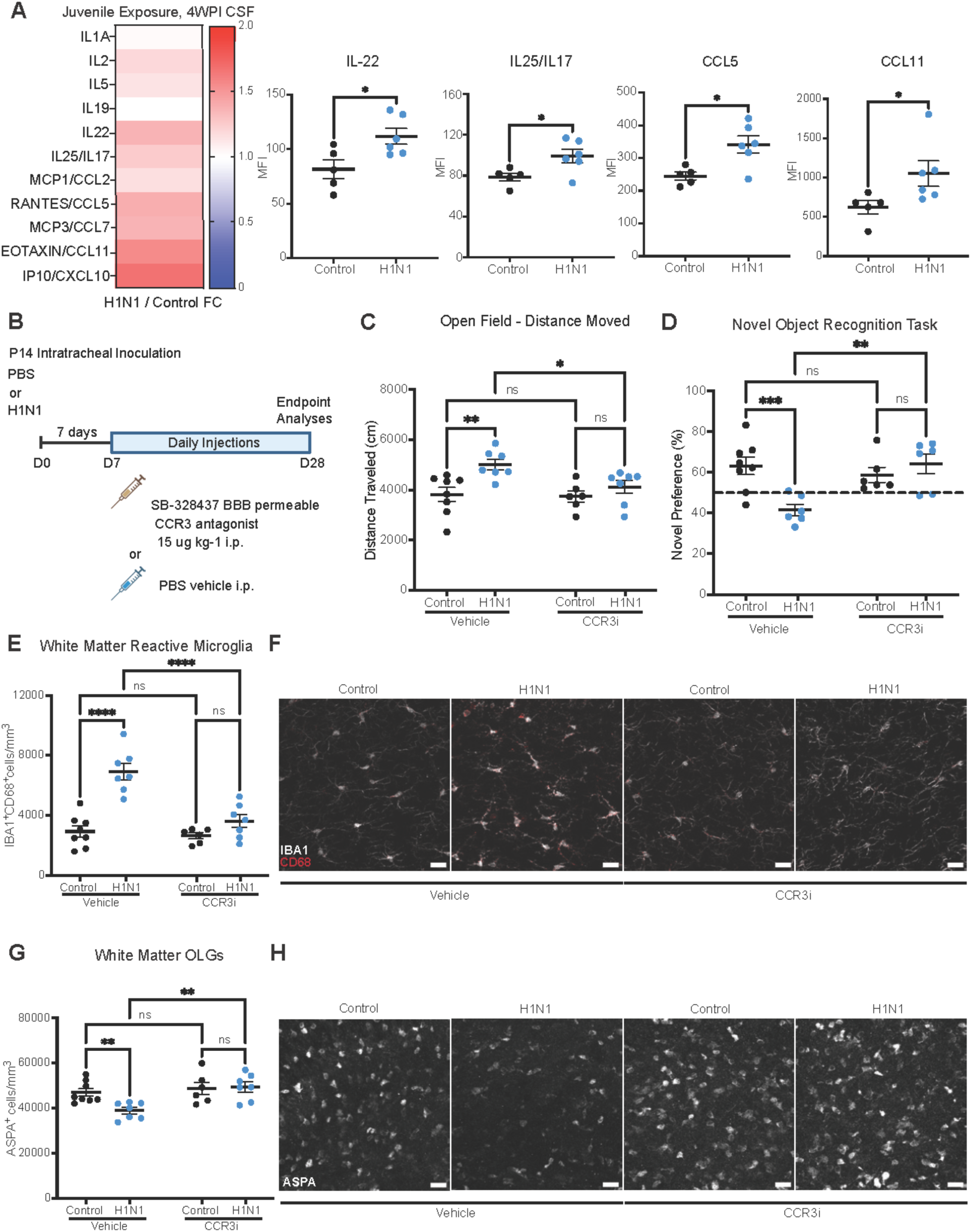
Chemokine receptor inhibition rescues oligodendroglial dysregulation and cognitive performance in novel object recognition testing. (A) Left, heatmap of fold change of CSF cytokine and chemokine levels in H1N1 infected mice relative to control mice, 4 weeks post-infection. Right, quantification of raw mean fluorescence intensity (MFI) of CSF levels of cytokines. *n* = 5 control, *n* = 6 H1N1 mice. (B) Experimental timeline with CCR3i treatment, all quantifications at 4 weeks post-infection (WPI). Mice were intratracheally inoculated with PBS or H1N1 at P14 and administered daily i.p. injections of vehicle or CCR3i for three weeks starting from P21 to P42. (C) Total distance moved in the open field over 10-minute testing period. *n* = 8, 7, 6, and 7 mice per group. (D) Novel preference for NORT. *n* = 8, 7, 6, and 7 mice per group. (E) Quantification of white matter reactive microglia (IBA1^+^CD68^+^) in the cingulum. *n* = 8, 7, 6, and 7 mice per group. (F) Representative confocal micrographs of white matter reactive microglia (IBA1, white; CD68, red) in the cingulum. (G) Quantification of white matter mature oligodendrocytes (ASPA^+^) in the cingulum. *n* = 8, 7, 6, and 7 mice per group. (H) Representative confocal micrographs of white matter mature oligodendrocytes (ASPA, white) in the cingulum. Data shown as mean ± SEM; each dot represents an individual mouse. ns: p > 0.05, * p < 0.05, ** p < 0.01, *** p < 0.001, **** p < 0.0001, analyzed via (B-D) unpaired two-tailed t-test or (F, G, H, and J) two-way ANOVA. (I, K) Scale bars, 40 μm.

With CCR3 inhibition, there continued to be no differences in the overall number of microglia across all groups in the cortex, subcortical white matter, and hippocampal white matter (**Figures S8D-S8F**). Concordant with the rescue of motor activity and attentional behavioral aberrations described above, CCR3 inhibition normalized the number of reactive microglia and mature oligodendrocytes in the subcortical white matter following juvenile H1N1 infection (**Figures 6E-6H**). However, CCR3 inhibition did not reduce the increased microglial reactivity found in the hippocampal white matter (**Figures S8G and S8J**), concordant with the lack of T-maze performance rescue. Reactive microglia in the cortex remained unchanged (**Figure S8H**), as did the number of OPCs in the subcortical white matter (**Figure S8I**). Taken together, these findings suggest that targeting CCR3-mediated chemokine signaling can partially mitigate the neurobiological effects of juvenile H1N1 infection by modulating microglial reactivity and oligodendrocyte integrity in subcortical white matter.

## DISCUSSION

Persistent cognitive and neuropsychiatric symptoms are commonly described after inflammatory illnesses, including respiratory infections^55,56^. Post-acute infection syndromes (PAIS) have gained more attention in recent years in the context of the COVID-19 pandemic. Although persistent cognitive impairment following COVID-19 infection is now well-recognized as a relatively common sequela in adults, children are also experiencing Long COVID neurological symptoms^48–50,57,58^. Currently, over 6 million children in the United States are living with Long COVID, with up to 44% experiencing cognitive and neuropsychiatric symptoms such as difficulty concentrating, memory problems, anxiety, and depression^48^. These neurological sequelae can manifest differently in children compared to adults, presenting as hyperactivity, decreased school performance, and behavioral problems^48,57^. Every year, millions of children worldwide contract influenza, with thousands requiring hospitalization, and children under the age of five are most severely affected^59–61^. This age group is especially vulnerable to respiratory infections, such as influenza and respiratory syncytial virus (RSV)^62–66^. Many survivors of the 1918 influenza pandemic experienced lasting neuropsychiatric symptoms^67–69^, and impairments in attention and other cognitive domains have been well-documented in people following non-pandemic influenza (for review, see Smith et al.^33^).

Taken together, the findings presented reveal that a relatively mild, non-neurotropic respiratory H1N1 influenza infection during the juvenile developmental period is sufficient to induce white matter-selective microglial reactivity, perturb oligodendroglial lineage dynamics, impair myelin development, and produce lasting cognitive changes in mice. The developmental selectivity we observe, lasting effects after H1N1 influenza infection during the juvenile but not the adult period, highlights neural-immune vulnerability during early-life periods of active circuit remodeling and myelination. This is consistent with the concept that systemic inflammation can drive a white matter-selective microglial response and resulting disruption of oligodendroglial homeostasis and plasticity, as has been demonstrated following multiple immune challenges in adults^1–4^, that is particularly consequential in the developing juvenile brain.

By one week after juvenile H1N1 influenza respiratory infection, microglia in subcortical and hippocampal white matter exhibit a reactive state without changes in total number, while mature oligodendrocytes are selectively reduced in subcortical white matter. Single-nucleus RNA sequencing places this early response in a mechanistic context: microglia upregulate innate immune and antigen-processing programs, including MHC I genes, and express chemokine, inflammatory, and interferon-responsive gene modules. Concomitantly, oligodendroglial composition shifts toward less mature states with induction of a disease-associated oligodendrocyte signature in mature cells. These convergent cell-state changes suggest a coordinated inflammatory milieu that could both delay or prevent oligodendrocyte maturation and shift mature oligodendrocytes into a stress-associated state, thereby reducing support for axons and impairing oligodendrogenesis during a period of active myelin development.

The functional implications of this juvenile neural-immune response evolve over time. Four weeks after infection, mice exhibit hyperlocomotion and impaired performance on an attention-dependent task, with persistent microglial reactivity and reduced numbers of mature oligodendrocytes in subcortical white matter. This behavioral phenotype of increased motor activity and decreased attentional function in mice is notable in the context of clinical reports of increased incidence of ADHD in children following COVID-19^70,71^, which can similarly cause persistent white matter-selective microglial reactivity in both mouse and human brains^2^. Whether respiratory COVID-19 infection in the juvenile period disrupts developmental myelination and causes a similar pattern of ADHD-like behavior in experimental models or in patients remains to be studied. Furthermore, we found that the elevated motor activity resolves with time, while increased anxiety-related behaviors emerge in later adulthood following juvenile respiratory immune challenge. These findings underscore the lasting and evolving consequences of even transient neuroinflammation and disrupted neurodevelopment. More broadly, these findings are consistent with emerging frameworks in which immune state is continuously integrated by the nervous system to shape behavior and cognition^72^.

It is important to note that four to eight weeks of ongoing neuroinflammation in a juvenile mouse represents a significant portion of the developmental myelination period for mice. In contrast, developmental myelination spans three decades in humans^21,22^. This context is important for interpretation: our data do not imply that a single, uncomplicated influenza infection in every child will lead to substantial disruption of myelin development or lasting cognitive and behavioral deficits. Rather, these findings imply that persistent or recurrent inflammatory states and the accumulation of immune insults over time could result in sustained microglial reactivity and impede oligodendroglial maturation, increasing the risk for long-lasting structural and functional changes. From this perspective, it is important to note that white matter microglial reactivity occurs following a wide range of systemic insults and is differentially persistent following different immune challenges^1–3^, for example lasting for longer periods of time following some respiratory viral infections such as COVID-19 compared to others such as H1N1 influenza^2^.

By eight weeks, microglial reactivity and mature oligodendrocyte number normalize in subcortical white matter following juvenile H1N1 influenza, yet cognitive deficits persist, and analysis of myelin ultrastructure reveals lasting reductions in myelinated axon density and myelin thickness, particularly in small- and medium-caliber axons that myelinate later^28,29^. This dissociation between cell number and myelin ultrastructure suggests that an early inflammatory derailment of developmental myelination can yield enduring structural deficits even after cell counts appear restored. Given the importance of myelin for conduction velocity, metabolic support, and neural circuit dynamics, coordination and function^15,73–76^, these alterations likely contribute to the sustained attention and short-term memory impairments observed here, which remain evident even six months after infection.

The chemokine axis appears to be a key driver and actionable link between systemic infection or other immune challenges^3,77,78^ and CNS changes. CSF chemokines including CCL11 and CCL5 are elevated, and microglia display elevated chemokine gene expression in single-nucleus sequencing analysis, similar to the chemokine-associated microglial phenotype identified after respiratory COVID-19^2^. Pharmacologic inhibition of the multi-chemokine receptor CCR3 during the subacute period after infection rescues many of the cellular and behavioral aberrations caused by juvenile H1N1 influenza infection, including normalizing hyperlocomotion, rescuing attention and memory in the Novel Object Recognition Task, and reducing microglial reactivity while restoring mature oligodendrocytes in subcortical white matter. These results identify CCR3 signaling as a tractable node in a broader chemokine-driven cascade linking peripheral infection to central white matter dysregulation and cognitive impairment. An open question, and a major focus of ongoing work, is the nature of the chemokine relay that transduces lung-localized respiratory infection into central neuroinflammation. Circulating cytokines and chemokines may enhance leukocyte and myeloid trafficking into the brain and its borders ^79^ (meninges, perivascular spaces, choroid plexus) and induce reactive states in resident immune cells. They may also stimulate microglia to produce additional chemokines and cytokines, establishing local feed-forward loops that sustain reactivity.

This work has several implications. Juvenile respiratory immune challenges can perturb developmental myelination with lasting cognitive and neuropsychiatric effects. In cases of persistent neuroinflammation, or repeated immune challenges during the childhood and adolescent period, neuroimmune-modulating strategies should be studied for at-risk pediatric populations. Prophylactic vaccination and intervening during or shortly after a major inflammatory insult may prevent long-term cellular and behavioral sequelae. Future studies should define the timing for effective intervention. The results presented here also indicate that the timing of immune challenges matter to the long-term neurological and neuropsychiatric consequences, with different possible consequences at different developmental stages.

### Limitations of the study

Several limitations of this study should be considered. While our data strongly implicate microglial reactivity as a central regulator of white matter dysregulation following juvenile influenza infection, we did not directly test the necessity of microglia in driving the observed oligodendroglial, myelin, and cognitive phenotypes. In previous studies of mouse models of cancer chemotherapy-^1,80^ and immunotherapy-related^3^ cognitive impairment, interventions that deplete microglia normalize multicellular dysfunction and improve cognitive outcomes. Given the striking similarities across various immune challenges in causing white matter-selective microglial reactivity and consequent dysregulation of oligodendrocytes and myelin, it is possible that analogous microglia-targeted strategies could mitigate cognitive impairment following major respiratory immune challenges. Genetic or pharmacologic microglial depletion approaches would be required to establish causal relationships between microglial reactivity and long-term cellular and behavioral outcomes following influenza.

In addition to their role in supporting oligodendrocyte and myelin homeostasis, microglia play a critical role in circuit refinement and synaptic pruning during normal postnatal brain development^81^. Microglia help sculpt neural circuits during sensitive developmental windows^82–85^, and disruptions to these processes have been linked to long-lasting alterations in circuit function and behavior^86–88^. It is likely that influenza-induced microglial reactivity also perturbs microglia-mediated synaptic remodeling, representing an additional mechanism by which juvenile immune challenges could produce enduring cognitive and neuropsychiatric consequences. Direct investigation of whether synaptic structure is altered following juvenile influenza and of how microglia-synapse interactions are perturbed, will be an important direction for future work.

Neurotoxic astrocytes are known to emerge downstream of reactive microglia and can contribute to oligodendrocyte toxicity and impaired synaptic support ^10,89^. Whether astrocytes adopt neurotoxic or maladaptive states following juvenile influenza infection, and whether such astrocyte responses contribute to the persistent myelin ultrastructural abnormalities observed here, remain important open questions.

Our work focuses on infection in the absence of vaccination or prior immune priming. Influenza vaccination in pediatric populations may significantly reduce viral replication, suppress inflammatory responses, and lower chemokine levels, thereby attenuating downstream consequences of infection. Whether vaccination attenuates or prevents the white matter-specific microglial reactivity and oligodendroglial disruption observed is an important translational question that warrants further study. Similarly, we did not examine the effects of repeated infections or combined immune challenges, which may more closely reflect real-world pediatric exposure and could exacerbate or prolong neural-immune sensitization.

While our data identify CCR3 signaling as a therapeutic target linking peripheral infection to central white matter pathology, the upstream chemokine relay that conveys lung-localized inflammation to the brain remains incompletely defined. The relative contributions of circulating cytokines and chemokines, leukocyte trafficking at CNS border sites, and feed-forward signaling within the brain require further investigation. Understanding these pathways will be critical for identifying points of intervention and determining whether targeting peripheral immune signaling, CNS-resident immune cells, or their interaction is most effective in preserving developmental myelination and cognitive function.

## CONCLUSIONS

These findings position major respiratory immune challenges during the juvenile period as a potential driver of durable white matter vulnerability and cognitive dysfunction. Our data indicate that immune activation during sensitive developmental windows can have consequences that extend well beyond the resolution of the illness. Understanding the neural-immune mechanisms by which transient systemic inflammation reshapes microglial state, oligodendroglial maturation, and myelin ultrastructure is essential for understanding how common pediatric respiratory infections may contribute to persistent cognitive and neuropsychiatric symptoms. The identification of a chemokine-driven pathway linking peripheral infection to central white matter dysregulation highlights a tractable therapeutic entry point and suggests that timely neural-immune modulation may preserve developmental myelination and cognitive function. These insights may inform strategies to mitigate persistent “brain fog” syndromes across a wide range of infectious and inflammatory diseases.

## ACKNOWLEDGEMENTS

This authors gratefully acknowledge support from by the HHMI Emerging Pathogens Initiative, The Gatsby Initiative in Brain Development and Psychiatry from the Gatsby Charitable Foundation, Waxman Family Research Fund, and the National Institute of Neurological Disorders and Stroke (F31NS143332). Model schematics were created with BioRender.com. The authors thank the Human Immune Monitoring Core at Stanford.

## AUTHOR CONTRIBUTIONS

Conceptualization, methodology, validation, and visualization were performed by M.M., K.M., and C.B.; formal analysis by K.M. and K.S.; resources provided by M.M; investigation was performed by K.M., K.S., S.A., L.N., N.Z-G., A.R., B.Y., E.H.C., T.P., A.G.; writing – original draft by M.M. and K.M.; writing – review & editing by M.M., K.M. and A.I.; data curation by K.M.; supervised by M.M. and A.G.; and funding acquisition by M.M. M.M. conceived of the projects and supervised all aspects of the work.

## Declaration of Interests

M.M. holds equity in MapLight Therapeutics and Stellaromics Inc. Stanford University has filed an intellectual property application with M.M and A.C.G. as inventors entitled “Chemokine receptor blockade for treatment of cognitive dysfunction following cancer immunotherapy”.

## Data Availability

Single nucleus RNA-sequencing are available on GEO with accession number GSE316351 (token orazycoctpsjpkv).

## METHODS

### EXPERIMENTAL METHODS AND SUBJECT DETAILS

#### Mouse Models

All procedures relating to animal care and treatment adhered to the Institutional Animal Care and Use Committee protocol at Stanford University and followed the guidelines by the National Institutes of Health. For experiments involving H1N1 influenza infection at postnatal day (P)14, embryonic day (E)15-16 CD1 dams were obtained from Charles River Laboratories. In experiments involving mice inoculated at P14, at least three litters per experiment were used. CD1 mice infected at 2 months old were also obtained from Charles River. For conditional deletion of *Myrf*, hemizygous *Pdgfra-CreER^TM^* (Jackason Laboratory, 018280) mice were bred with homozygous *Myrf^lox/lox^* mice (Jackson Laboratory, 010607), in order to generate hemizygous *Pdgfra-CreER^TM^* and heterozygous *Myrf^lox/+^* mice, which were backcrossed to homozygous *Myrf^lox/lox^* to achieve hemizygous *Pdfgra-CreER^TM^* and homozygous *Myrf^OPC-/-^* (*Myrf^lox/lox^*;*Pdgfra-CreER^TM^*). Littermates that did not have the *Pdgfra-CreER^TM^* were used as control animals. To initiate Cre-dependent deletion of *Myrf*, mice were intraperitoneally injected with 100 mg kg^-1^ tamoxifen (Sigma-Aldrich) for 4 consecutive days at P14. Both male and female mice were included in all experimental groups. Mice were housed at Stanford University under standard conditions, with 12h light: 12h dark cycle and ad libitum access to food and water. The specific age of the mice used in each experiment is detailed in the corresponding figures and text.

#### H1N1 influenza respiratory infection model

Gradient-purified influenza A/PR8/34 (H1N1) was obtained commercially from AVS Bio and stored in 5μL aliquots at -80°C. Mice were infected intratracheally with H1N1 as previously described^90^. Briefly, mice were anesthetized using 30% v/v isoflurane diluted in propylene glycol. A pipette was used to administer 20 μL of H1N1 in 1x phosphate buffered saline (PBS) intratracheally, with a dose of 1 PFU for P14 mice and 50 PFU for 2-month-old mice. Intratracheal administration of PBS was used as a control. Mice were weighed and monitored daily for two weeks post-inoculation to assess sickness behavior.

#### CCR3 inhibitor administration

P14 mice were infected with H1N1 influenza as described above. Seven days post-infection, animals were weighed and administered daily intraperitoneally injections of either the CCR3 inhibitor SB-328437 (15μg/kg; MedChem Express) or vehicle control, continuing until the study endpoint at day (D)28. Following treatment, animals underwent cognitive assays and CSF and brain collection as described below.

### METHOD DETAILS

#### CSF collection and perfusion

At the conclusion of each experiment, mice were anesthetized with isoflurane and positioned onto a stereotactic surgical rig (Stoelting,) with the head secured at a 30-degree upward angle. An incision was made through the skin and muscle, and the neck muscles were gently separated using blunt metal forceps to expose the cisterna magna. Excess blood was carefully cleaned using cotton tips until the area was clear. CSF was collected from the cisterna magna using a pulled clear capillary glass tube (Harvard Apparatus 30-0062, 1.5 OD x 1.17 ID x 100mm). The CSF was stored in a low-binding microcentrifuge tubes (Costar, 3206), frozen on dry ice, and kept at -80°C until assayed. Following CSF collection, mice were transcardially perfused with 20mL of 0.1M PBS. Brains were fixed in 4% PFA (paraformaldehyde) overnight at 4°C, then transferred to 30% sucrose solution for cryoprotection. Brains were embedded in Tissue-Tek (Sakura) and sectioned coronally at 40μm using a sliding microtome (Microm HM450; Thermo Scientific).

#### Immunohistochemistry

Free-floating coronal sections from a 1-in-6 series were washed three times in 1X TBS before being incubated in a blocking solution consisting of 3% normal donkey serum and 0.3% Triton X-100 in TBS at room temperature for one hour. Primary antibodies were diluted in 1% normal donkey serum with 0.3% Triton X-100 in TBS and incubated with the sections overnight at 4°C. The primary antibodies used were: chicken anti-IBA1 (1:1000, Synaptic Systems, 234 009), rat anti-CD68 (1:200, Abcam, ab53444), rabbit anti-ASPA (1:250, EMD Millipore, ABN1698), and goat anti-PDGFRα (1:500, R&D Systems, AF1062). The following day, sections were rinsed three times in 1X TBS and incubated in a secondary antibody solution at room temperature, protected from light, for two hours. The secondary antibodies included: Alexa 647 donkey anti-chicken IgG (1:500, Jackson Immunoresearch), Alexa 488 donkey anti-rat IgG (1:500, Jackson Immunoresearch), Alexa 594 donkey anti-goat IgG (1:500, Jackson Immunoresearch), and Alexa 555 donkey anti-rabbit IgG (1:500, Jackson Immunoresearch), all diluted in 1% blocking solution. Sections were then rinsed three times in 1X TBS, incubated with DAPI (1:1000, Thermo Fisher Scientific) for 5 minutes, and mounted using ProLong Gold mounting medium (Life Technologies).

#### Confocal imaging and quantification

Cell counting was performed by experimenters blinded to experimental conditions. Images were acquired using a Zeiss LSM980 scanning confocal microscope (Zeiss) at 20X magnification and analyzed using QuPath software. For microglia imaging, three consecutive sections were selected from the cortex and corpus callosum, approximately spanning bregma +1.2 to +0.8, and four consecutive sections were selected from the hilus of the dentate gyrus, approximately spanning bregma -1.8 to -2.4. For each section, the deep cortex, cingulum of the corpus callosum, and hilus of the dentate gyrus were identified, and two 424.3 x 424.3 μm fields per slice were selected in those areas for quantification. CD68^+^ and IBA1^+^ cells were counted manually in those regions. For the analysis of oligodendrocytes (ASPA^+^) and oligodendrocyte precursor cells (PDGFRα^+^), three consecutive sections spanning bregma +1.2 to +0.8 were stained. The cingulum in each hemisphere was imaged, with two 424.3 x 424.3 μm fields selected for manual quantification.

#### Transmission Electron Microscopy

Eight weeks post-infection (P70 for mice infected during the juvenile period and P116 for mice infected during adulthood) or 6 months post-infection (P194 for mice infected during the juvenile period), mice were anesthetized with isoflurane. Mice were then sacrificed via transcardial perfusion using Karnovsky’s fixative, consisting of 2% glutaraldehyde (Electron Microscopy Sciences, EMS 16000) and 4% PFA (EMS 15700) in 0.1 M sodium cacodylate (EMS 12300), pH 7. Transmission electron microscopy (TEM) was performed on the premotor cortex (M2) subcortical fibers as they exit cortical layer VI and enter the corpus callosum. Samples were post-fixed in 1% osmium tetroxide (EMS 19100) for 1 hour at room temperature, washed three times with ultrafiltered water, and stained en bloc for 2 hours at room temperature. Samples were dehydrated in graded ethanol solutions (50%, 75%, and 95%) for 15 minutes each at 4°C, equilibrated to room temperature, and rinsed twice in 100% ethanol, followed by acetonitrile for 15 minutes. Samples were infiltrated with EMbed-812 resin (EMS 14120), initially mixed 1:1 with acetonitrile for 2 hours, followed by a 2:1 EMbed-812: acetonitrile mixture for another 2 hours. Subsequently, samples were placed in pure EMbed-812 for 2 hours, transferred to TAAB capsules filled with fresh resin, and cured in a 65°C oven overnight. Sections were cut between 75 and 90 nm using a Leica Ultracut S (Leica, Wetzlar, Germany) and mounted on Formvar/carbon-coated slot grids (EMS FCF2010-Cu) or 100 mesh Cu grids (EMS FCF100-Cu). Grids were contrast-stained for 30 seconds in 3.5% uranyl acetate in 50% acetone, followed by 30 seconds in 0.2% lead citrate. Imaging was performed using a JEOL JEM-1400 TEM at 120 kV, with images captured using a Gatan Orius digital camera. Image analyses were conducted by experimenters blinded to experimental conditions. Axons in the cingulum of the corpus callosum were analyzed for *g*-ratios, calculated by dividing the shortest axonal diameter by the corresponding axon-plus-sheath diameter (diameter of axon/diameter of axon plus myelin sheath), as well as for myelinated axon density (total number of myelinated axons per frame). A minimum of 1,500 axons were scored per group. Myelinated axon density was determined by quantifying the number of myelinated axons per 6,000X electron micrograph, with an average of 23 images quantified per mouse. The total number of myelinated axons was divided by the total area quantified per animal. Approximately, 150-400 axons were scored for each mouse. Group means were calculated on a per mouse basis, not per axon or image.

#### Mouse CSF cytokine analysis

The Luminex multiplex assay was conducted by the Human Immune Monitoring Center at Stanford University. Mouse 48-plex Procarta kits were obtained from Thermo Fisher (Santa Clara, California, USA) and used according to the manufacturer’s instructions, with some modifications as detailed below. Briefly, beads were added to a 96-well plate and washed using a Biotek Elx405 washer. Samples were then added to the plate containing the mixed antibody-linked beads and incubated overnight at 4°C with shaking. Both cold (4°C) and room temperature incubation steps were performed on an orbital shaker set to 500-600 RPM. Following the overnight incubation, the plates were washed again using the Biotek Elx405 washer. A biotinylated detection antibody was added and incubated for 60 minutes at room temperature with shaking. The plates were washed as previously described, and streptavidin-PE was added. After a 30-minute incubation at room temperature, the plates were washed again, and reading buffer was added to the wells. CSF samples were measured as singlets, while serum samples were measured in duplicate. Plates were read on an FM3D FlexMap instrument, ensuring a minimum of 50 beads per sample per cytokine. Custom Assay Chex control beads, purchased from Radix Biosolutions (Georgetown, Texas), were added to all wells. The assay was conducted in triplicates. For each cytokine, we first calculated the mean of the mean fluorescent intensities (MFIs) for the control group. To determine the fold changes (FCs) for each cytokine, we divided the MFI of each individual influenza sample by the mean MFI of the control group. We then averaged these FCs to obtain a single representative value for each cytokine, which was included in the heat map. For plots displaying individual cytokines, the MFIs for each animal were plotted, and statistical analyses were performed on a per-animal basis.

#### Open Field Test

To evaluate anxiety-like behavior and locomotion, mice were placed in a square, opaque enclosure measuring 50 cm x 50 cm with 50 cm high walls and allowed to explore freely for 10 minutes. Prior to the open field test, mice were handled for 5 minutes daily over two days. The test was conducted the first time the mice were placed in the enclosure to assess baseline behaviors before habituation to the experimental chamber. Testing was conducted during the animals’ rest phase in a dark room illuminated only by red light.

Behaviors, including distance moved and velocity, were recorded with a camera mounted 115 cm above the chamber and analyzed using EthoVision video tracking software (Noldus). Time spent in the center of the enclosure was measured, defined as time spent at least 10 cm away from the walls. After testing each subject, the entire chamber was cleaned with 70% ethanol. The number of fecal boli in the chamber was counted at the end of the 10-minute period.

#### Novel Object Recognition Task

Cognition was assessed using a modified version of the Novel Object Recognition Task (NORT) that emphasized the attentional component of the task. The test was adjusted to shorten the interval between the training and testing phases, thereby increasing the cognitive load on short-term memory and attention (<5 min), instead of longer-term memory and hippocampal function. The timing of NORT testing is detailed in each figure. Animals were handled for 5 minutes daily over the 5 days preceding the test. Following handling, mice were placed in the experimental chamber for 10 minutes daily over the 3 days preceding the test. The setup consisted of an opaque Plexiglas chamber measuring 50 cm x 50 cm x 50 cm, with a camera mounted 115 cm above the chamber. Testing was conducted during the animals’ rest phase in a dark room illuminated only by red light. On the testing day, mice were brought to the testing room and allowed to settle in their cages for 30 minutes before handling. After handling for 2 minutes, mice were acclimated in the chamber for 10 minutes, then returned to their home cage for 5 minutes. Mice were then placed in the experimental chamber with two identical Lego objects, each approximately 4 cm in size. Each time a mouse was placed into the chamber, it faced the opaque wall with its tail directed towards the objects. During the training phase, mice explored the two identical objects for 5 minutes before being returned to their home cage for 5 minutes while the chamber and objects were cleaned with 70% ethanol. For the novel object phase, one cleaned object from the sample phase was placed back into the chamber along with a new Lego object of similar size. During the novel object testing phase, mice were allowed to explore for 10 minutes. The objects used as novel and familiar were counterbalanced, as was the position of the novel object across trials and animals. All Lego objects used in the behavioral paradigm were piloted to ensure no bias or object preference among the animals. Camera footage was analyzed using Noldus analysis software. Exploratory head gestures within 2 cm of the Lego object, including sniffing and biting, were considered object investigation, whereas sitting on the object or casual touching in passing was not. Only animals that explored both identical objects during the testing phase for a minimum of 20 seconds were included in the analysis. The Novel Preference was calculated as the percentage of time spent investigating the novel object relative to the total time spent investigating both objects: (time spent with novel object) / (time spent with novel object + time spent with familiar object).

#### Spontaneous Alternation T Maze Task

Hippocampal-dependent cognitive performance was assessed using the Spontaneous Alternation T Maze Task, which evaluates spatial working memory. This test was administered over two consecutive days, beginning immediately after the completion of the NORT assessment. The timing of Spontaneous Alternation T Maze testing is detailed in each figure. By this stage, mice had been handled prior to and during the NORT. Testing was conducted during the animals’ rest phase in a dark room illuminated only by red light. The T maze setup consisted of a start arm and two goal arms, each measuring 30 x 10 cm with 20 cm high walls. One goal arm was textured to allow for clear distinction by the mouse. On day 1 of the T maze assessment, following the completion of NORT and a minimum of 2 hours of rest, each mouse was placed at the base of a T-shaped maze facing away from the goal arms. As the mouse explored the maze, it chose an arm by entering it with all four paws. Upon arm selection, a guillotine door was closed, allowing the mouse to explore and familiarize itself with the chosen arm for 30 seconds. The mouse was then returned to the base of the maze facing away from the goal arms. This process was repeated for 11 consecutive cycles, allowing for 10 total choices. Animals that failed to alternate in over 50% of these trials were excluded from further analysis. On day 2, mice were brought into the testing room and allowed to settle in their cages for 30 minutes before handling. Each mouse was then handled for 2 minutes prior to T maze testing. Subsequently, a mouse was placed at the base of the T-shaped maze facing away from the goal arms. The mouse chose an arm by entering it with all four paws, at which point the guillotine door was closed to allow exploration of the chosen arm for 30 seconds. The mouse was then returned to its home cage for 15 minutes. After this interval, the mouse was placed back at the base of the maze facing away from the goal arms and allowed to explore and choose a goal arm. This process was repeated for 11 consecutive cycles. The chosen arms were recorded during testing, and mice were scored based on their performance over 10 trials.

#### Three Chamber Sociability Test

To assess sociability, defined as the preference to investigate a social versus a non-social stimulus, we employed a three-chambered sociability test. This test utilized a three-chambered arena with openings that allowed passage between the chambers. The three-chamber acrylic experimental cage (each chamber measuring 45 x 20 x 25 cm) contained two metal grid pencil cups (10 cm in diameter) in each of the outer chambers. The chambers were divided by transparent acrylic walls with 15-cm-wide entrance holes. One chamber contained a novel same-sex juvenile mouse in the pencil cup, while the other chamber contained an inanimate object in the pencil cup. Testing was conducted during the animals’ rest phase in a dark room illuminated only by red light. Subject animals were placed in the middle chamber and allowed to freely investigate each stimulus over a 10-minute period. Mice were tested for sociability one day after the conclusion of day 2 of the Spontaneous Alternation T Maze Task. Animals were habituated to the experimental room and the three-chamber experimental cage for 10 minutes one day prior to testing. The test mouse was initially placed in the middle chamber, and 1 minute later, the chamber dividers were removed, allowing the mouse to explore each chamber freely for 10 minutes. After testing each subject, the entire chamber was cleaned with 70% ethanol. The locations of the novel mouse and the inanimate object were counterbalanced between sessions. The time spent interacting with each pencil cup was analyzed using Ethovision XT software (Noldus) in an automated and condition-blinded manner. Social preference was calculated as the percentage of time spent investigating the novel mouse relative to the total time spent investigating both objects: (time spent with novel mouse) / (time spent with novel mouse + time spent with the inanimate object).

#### Brain Tissue Processing for Single-Nucleus RNA Sequencing

Juvenile mice (PBS control, *n* = 3; H1N1 PFU, *n* = 3) were intratracheally inoculated with PBS or H1N1 at P14 as described above. At 7 days post-infection, both PBS (*n* = 3) and H1N1 (*n* = 3) mice were transcardially perfused with 20 mL of perfusion buffer (110 mM NaCl, 10 mM HEPES, 25 mM glucose, 75 mM MgCl_2_, 2.5 mM KCl, 5 μg/mL^-1^ actinomycin D, and 10 μM triptolide in UltraPure DNase/RNase-free distilled water). The brains were quickly collected, flash frozen in isopentane on dry ice, and stored at -80°C. Frozen brains were sectioned inside a cryostat until the anterior-posterior (AP) coordinate reached 1.15 mm, and the cingulum from both hemispheres was punched out using a 1 mm biopsy punch with approximately 2 mm depth.

#### Single Nuclei Isolation and Sorting

Brain punches from each mouse were added to a Wheaton Dounce homogenizer containing 1 mL of ice-cold nuclei isolation medium (10 mM Tris pH 8.0, 250 mM sucrose, 25 mM KCl, 5 mM MgCl2, 0.1% Triton X-100, 1% RNasin Plus, 1x protease inhibitor, 0.1 mM DTT, 5 μg/mL^-1^ actinomycin D, 10 μM triptolide, and 10 μM anisomycin in UltraPure DNase/RNase-free distilled water). The tissue was dissociated by performing ten strokes with the loose Dounce pestle followed by ten strokes with the tight pestle. Homogenates were passed through a 40 μm cell strainer and centrifuged at 900g for 15 minutes to pellet the nuclei. The nuclei were resuspended in 1 mL of resuspension buffer (1x PBS, 1% nuclease-free BSA, and 0.5% RNasin Plus) and centrifuged again at 900g for 15 minutes. The supernatant was removed, and the nuclei were stained by adding 500 μL of resuspension buffer containing 0.1 μg/mL^-1^ DAPI and transferred to fluorescence-activated cell sorting (FACS) tubes. Nuclei were then sorted using a Sony MA900 sorter with a 70 μm chip. A standard gating strategy was applied to all samples, first gating nuclei based on size and scatter properties, then using doublet discrimination gates to exclude aggregates, and finally gating on DAPI. Single nuclei were sorted into chilled PCR tubes containing 10 μL of resuspension buffer. Nuclei counts were verified using a Fuchs-Rosenthal counting chamber.

#### RNA-seq Library Construction and Sequencing

RNA-seq libraries were prepared using the 10X Chromium Next GEM Single Cell 3’ ST Kit v3.1, following the manufacturer’s instructions. Nuclei were mixed with the master mix and loaded onto a 10X Chromium Next GEM Chip G, aiming to recover 10,000 cells from each sample. Libraries were generated according to the manufacturer’s protocols, with quality controls performed after complementary DNA amplification and library construction to ensure sample quality using Agilent 4200 TapeStation and High Sensitivity D5000 ScreenTape. Libraries were loaded at 650 pM along with 1% PhiX control on an Illumina NextSeq2000 and paired-end sequenced (28 cycles read1, 10 cycles i7 index, 10 cycles i5 index, 90 cycles read2) at a targeted depth of 20,000 reads per nucleus.

#### RNA-seq Data Analysis

Raw read (FASTQ) files were aligned to the mouse reference genome (mm10-2020-A) using Cell Ranger (v.7.2.0). For single-nucleus RNA sequencing, all analysis, quantification, and statistical testing were completed using R (v.4.2.0). Seurat (v.5.0.3) was used for preprocessing, dimensionality reduction, clustering, and differential expression testing. Cell Ranger count matrices for each sample were merged, and low-quality cells with fewer than 500 detected genes or greater than 3% mitochondrial mapping unique molecular identifiers (UMIs) and putative doublets with greater than 3,500 detected genes were removed. A total of 54,226 nuclei passed these quality control criteria (26,653 PBS and 27,5723 H1N1). UMI count data were normalized by dividing the counts per gene per cell by the total UMIs per cell, multiplying by a scale factor of 10,000, and applying natural log transformation. Variance stabilizing transformation was performed, and the top 2,000 variable genes were identified, followed by principal component analysis. UMAP was conducted using the first 20 principal components, and graph-based clustering identified clusters with a resolution parameter of 0.8. To focus on oligodendroglial lineage cells, oligodendrocytes and OPCs were subsetted and reclustered using similar parameters. Trajectories from OPCs to mature oligodendrocytes were labeled using previously defined markers for oligodendroglial subpopulations^53^. Differential expression testing was performed using Wilcoxon rank-sum tests with Bonferroni corrections for multiple comparisons. Adjusted p<0.05 with average log2-fold changes greater than 0.25 were considered statistically significant. ScType and differential expression testing identified cell types based on canonical marker gene expression, and clusters of the same cell type were merged. To produce violin plots, for each sample, we performed first cell-level normalization, and then centered the gene expression around 0 to allow PCA computation. Following the PCA reduction, we clustered the cells using shared nearest neighbor clustering. To examine the various state signatures of each of the nuclei in the microglia and mature oligodendrocyte cluster, we used the function AddModuleScore by the Seurat package, which calculates the average expression levels of the gene set subtracted by the aggregated expression of 100 randomly chosen control gene sets, where the control gene sets are chosen from matching 25 expression bins corresponding to the tested gene set expression. To calculate the *p* values, we pseudobulked the nuclei from each to sample to generate average gene expression levels per sample using AggregateExpression by the Seurat package. The *p* values were calculated using unpaired t-test. The microglial state scores (chemokine, inflammatory, and interferon response) were determined using previously defined markers for microglial substates^2^. Total chemokine state genes included: *Ccl4, Ccl3, Atf3, Cd83, Egfr1, Ccl12, Slc15a3, Csf1, Cxcl10, Id2, Stk38l, C5ar1, C3ar1, Plau, Plekho2, Hif1a, Eif4a1, Ccl9,* and *Fam20c*. Total inflammatory state genes included: *Cd83, H2-K1, Gpr84, Ccl12, Icam1, Tlr2, Nfkbia, Cxcl16, H2-D1, Tnfaip2, Slc15a3, Il1a, Hivep3, Marcksl1, Rassf4, Junb, Vcam1, Herpud1, Lacc1,* and *Cxcl10*. Total interferon response state genes included: *Ccl12, Cxcl10, Ifit3, Ccl5, Lgals3bp, Ifitm3, Slfn5, Ifit2, Bst2, Stat1, H2-K1, Parp14, H2-D1, Trim30a, Cxcl13, Rnf213, Ly6a, Ifi27l2a, Isg15,* and *Eif2ak2*. Disease-associated oligodendrocyte signature scores were determined using previously defined markers^54^ and included: *B2m, C4b, Cd63, Cd9, Clptm1, Ctsb, Fabp5, Gpd1, Gstp1, H2-K1, H2-D1, H3f3a, Ifi27, Il33, Klk6, Lbh, Mir7067, Opalin, Plekha1, Ptma, Rnase4, Rpl26, Rps2, Serpina3n, Sgk1,* and *Stmn1*. Gene set enrichment analysis was performed using Cluster Profiler (v.4.6.2) to identify enrichment of gene ontology biological process terms in cluster differentially expressed genes using a hypergeometric test^91^.

#### RNA Extraction, cDNA Synthesis, and qRT-PCR

Following perfusion, tissue samples were collected and flash frozen. To extract RNA, 600 μL of Buffer RLT was added to the tissue, which was then homogenized using the TissueLyser LT. The RNeasy® Plus Mini Kit (Qiagen) was employed for RNA extraction from the homogenized tissue. After homogenization, the lysate was centrifuged at full speed for 3 minutes, and the supernatant was removed. An equal volume of 70% ethanol was added to the supernatant and mixed by pipetting. The mixture was transferred to a RNeasy spin column and centrifuged for 15 seconds at 10,000 rpm, with the flow-through discarded. Next, 700 μL of Buffer RW1 was added to the spin column, centrifuged for 15 seconds at 10,000 rpm, and the flow-through discarded. This was followed by two washes with 500 μL of Buffer RPE: the first wash was centrifuged for 15 seconds at 10,000 rpm, and the second wash for 2 minutes at the same speed to remove residual ethanol. To elute the RNA, 30 μL of RNase-free water was added to the spin column membrane, and the sample was centrifuged for 1 minute at 10,000 rpm. For qRT-PCR, reactions were performed using a Mastercycler ep realplex (Eppendorf) with the Power SYBR® Green RNA-to-CT™ 1-Step Kit (Applied Biosystems). The master mix included Power SYBR® Green RT-PCR Mix, forward and reverse primers, RT enzyme mix, RNA template, and RNase-free water, which were pipetted into the wells of a reaction plate. The plate was sealed with optical adhesive film, briefly centrifuged, and run on the Mastercycler. All PCR primers were designed in-house and purchased from Integrated DNA Technologies. Relative gene expression was calculated using the 2-ΔΔCT method, normalized to the housekeeping gene 18S. Primer sequences are as follows: 18S, F: GAA TAA TGG AAT AGG ACC GC, R: CTT TCG CTC TGG TCC GTC TT; NP, F: GCC AGA ATG CCA CTG AAA , R: GAT CAA CCG TCC CTC ATA ATC.

### QUANTIFICATION AND STATISTICAL ANALYSIS

For all quantifications, experimenters were blinded to sample identity and condition. Fluorescent immunohistochemistry and confocal microscopy images were captured at 20X magnification, with cells considered co-labeled when markers co-localized within the same plane. For cortical and white matter microglia staining, three sections per mouse were analyzed, with four frames in standardized locations per section (totaling 12 images per mouse). For hippocampal microglia staining, four sections per mouse were analyzed, with two frames in standardized locations per section (totaling 8 images per mouse). For oligodendrocyte staining, three sections per mouse were analyzed, with four frames in standardized locations per section (totaling 12 images per mouse). Approximately 300 to 1,200 cells were counted per mouse for each immunohistochemical marker analysis. Cell density was calculated by dividing the total number of cells quantified for each lineage by the total volume of the imaged frames (mm³). In all experiments, “n” refers to the number of mice, with “n” representing at least 3 mice per group. The specific “n” for each experiment is detailed in the figure legends.

Statistical analyses were performed using Prism Software (GraphPad). For comparisons involving three or more comparison groups and one or two variables, one-way or two-way ANOVAs, respectively, were used with Tukey’s multiple comparisons post hoc corrections to assess main group differences. When stratifying by axon diameter, analyses were performed using unpaired two-tailed Student’s t-tests with correction for multiple comparisons. For analyses involving only two groups, unpaired two-tailed Student’s t-tests were employed. The Shapiro-Wilk test was used to determine normality for all datasets. The Mann-Whitney U test was used for datasets that were not normally distributed, as indicated in the figure legend. A significance level of p < 0.05 was used to designate significant differences.

---

**Figure S1.**
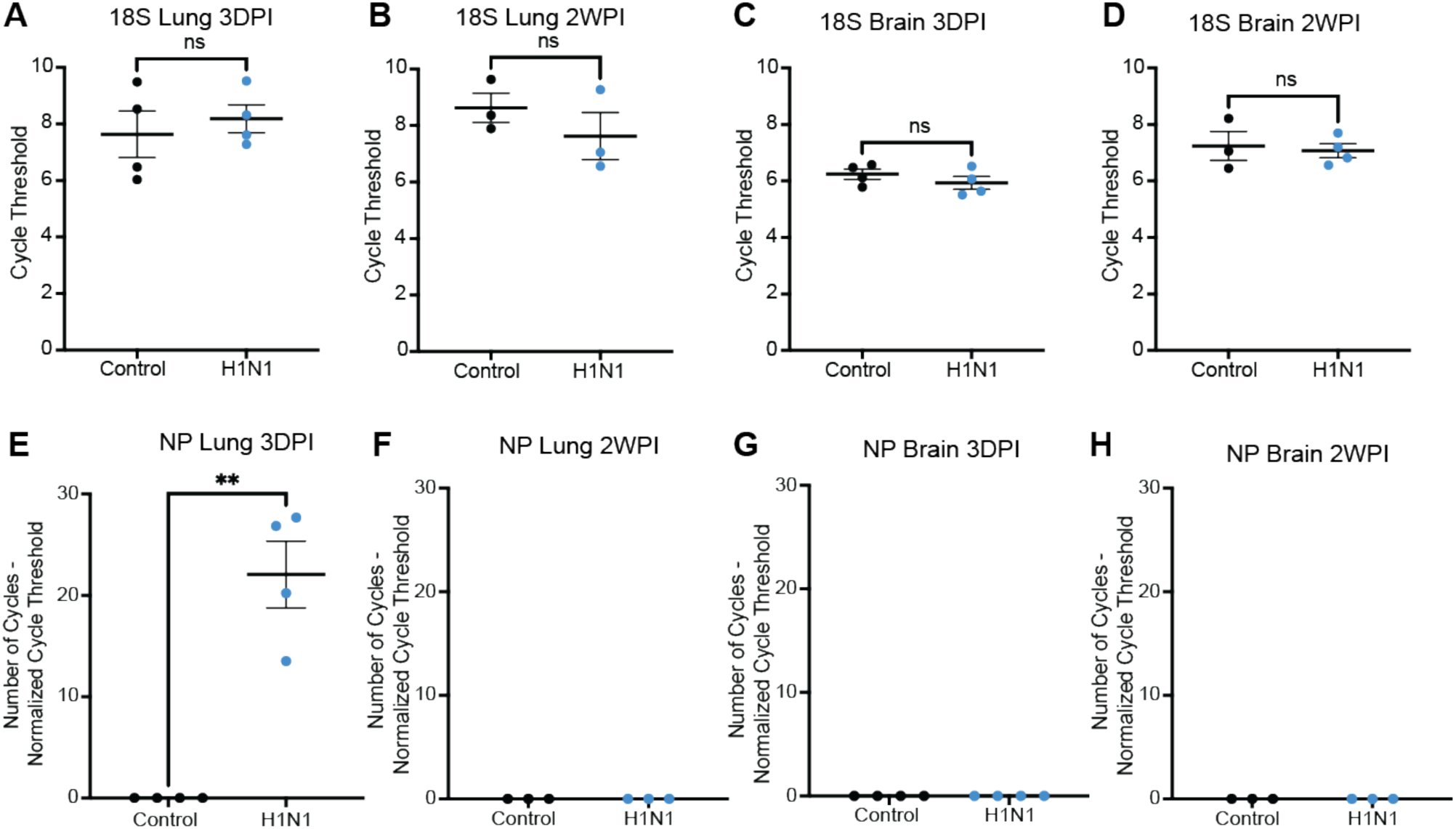
H1N1 virus is confined to the lung and cleared by two weeks, related to Figure 1. (A-D) Quantification of cycle threshold (CT) of 18S in the (A and B) lung or (C and D) brain at (A and C) 3 days (DPI) or (B and D) 2 weeks post-infection (WPI). (E-H) Quantification of influenza virus nucleoprotein (NP) normalized CT in the (E and F) lung or (G and H) brain at (E and G) 3 days or (F and H) 2 weeks post-infection. Data shown as mean ± SEM; each dot represents an individual mouse. ns: p > 0.05, ** p < 0.01, analyzed via unpaired two-tailed t-test.

**Figure S2.**
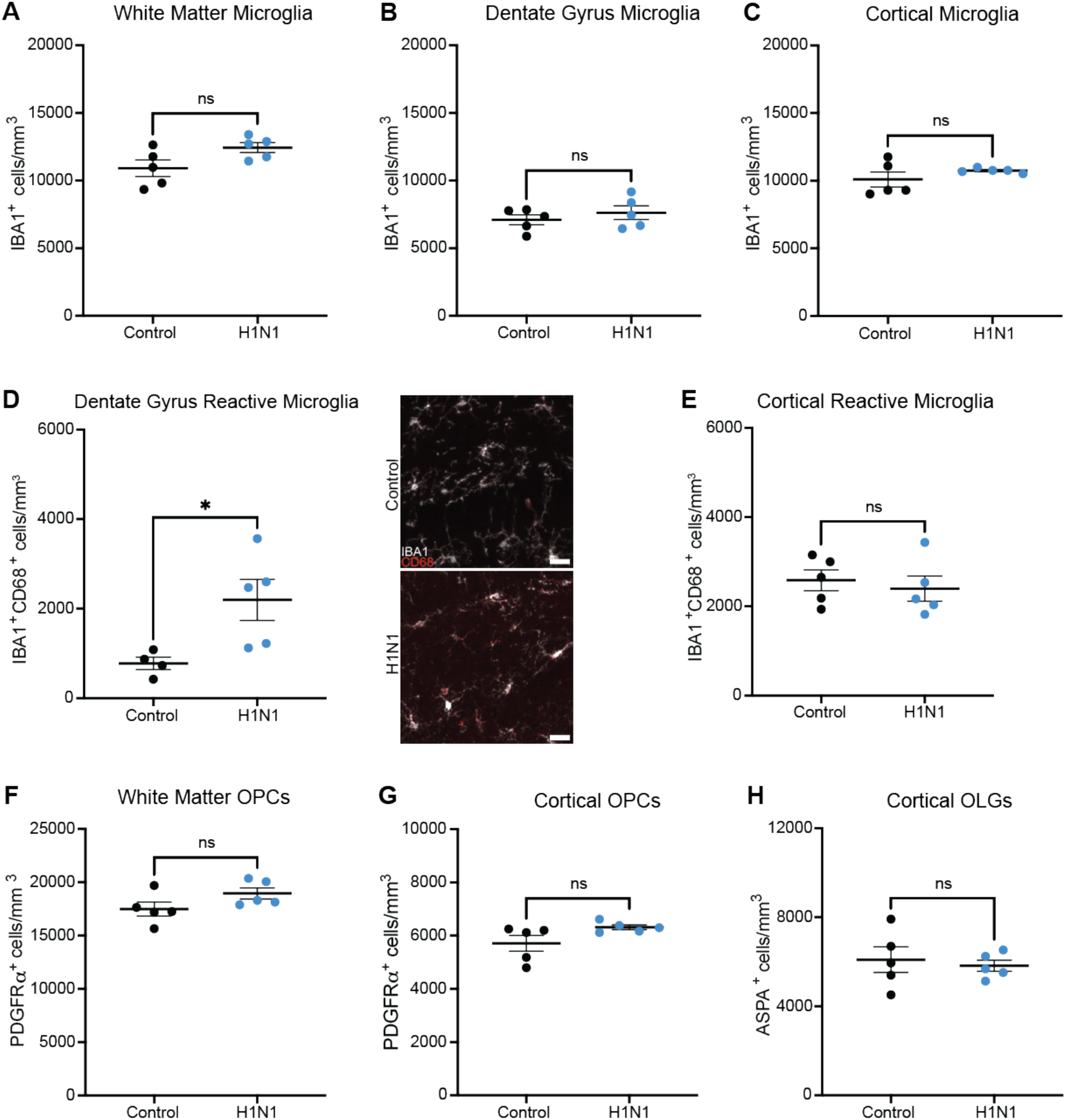
Glial cell counts 7 days post-infection in the white matter, dentate gyrus, and cortex, related to Figure 1. (A) Quantification of white matter microglia (IBA1^+^) 7 days post-infection in the cingulum. *n* = 5 mice per group. (B) Quantification of white matter microglia (IBA1^+^) 7 days post-infection in the hilus of the dentate gyrus. *n* = 5 mice per group. (C) Quantification of gray matter microglia (IBA1^+^) 7 days post-infection in the secondary motor cortex (M2). *n* = 5 mice per group. (D) Quantification and representative confocal micrographs of white matter reactive microglia (IBA1^+^CD68^+^) 7 days post-infection in the hilus of the dentate gyrus. *n* = 4 control, *n* = 5 H1N1 mice. (E) Quantification of gray matter reactive microglia (IBA1^+^ CD68^+^) 7 days post-infection in the M2 region of cortex. *n* = 5 mice per group. (F) Quantification of white matter OPCs (PDGFRα^+^) 7 days post-infection in the cingulum. *n* = 5 mice per group. (G) Quantification of gray matter OPCs (PDGFRα^+^) 7 days post-infection in the M2 region of cortex. *n* = 5 mice per group. (H) Quantification of gray matter mature oligodendrocytes (ASPA^+^) 7 days post-infection in the M2 region of cortex. *n* = 5 mice per group. Data shown as mean ± SEM; each dot represents an individual mouse. ns: p > 0.05, * p < 0.05, analyzed via unpaired two-tailed t-test. (D) Scale bars, 40 μm.

**Figure S3.**
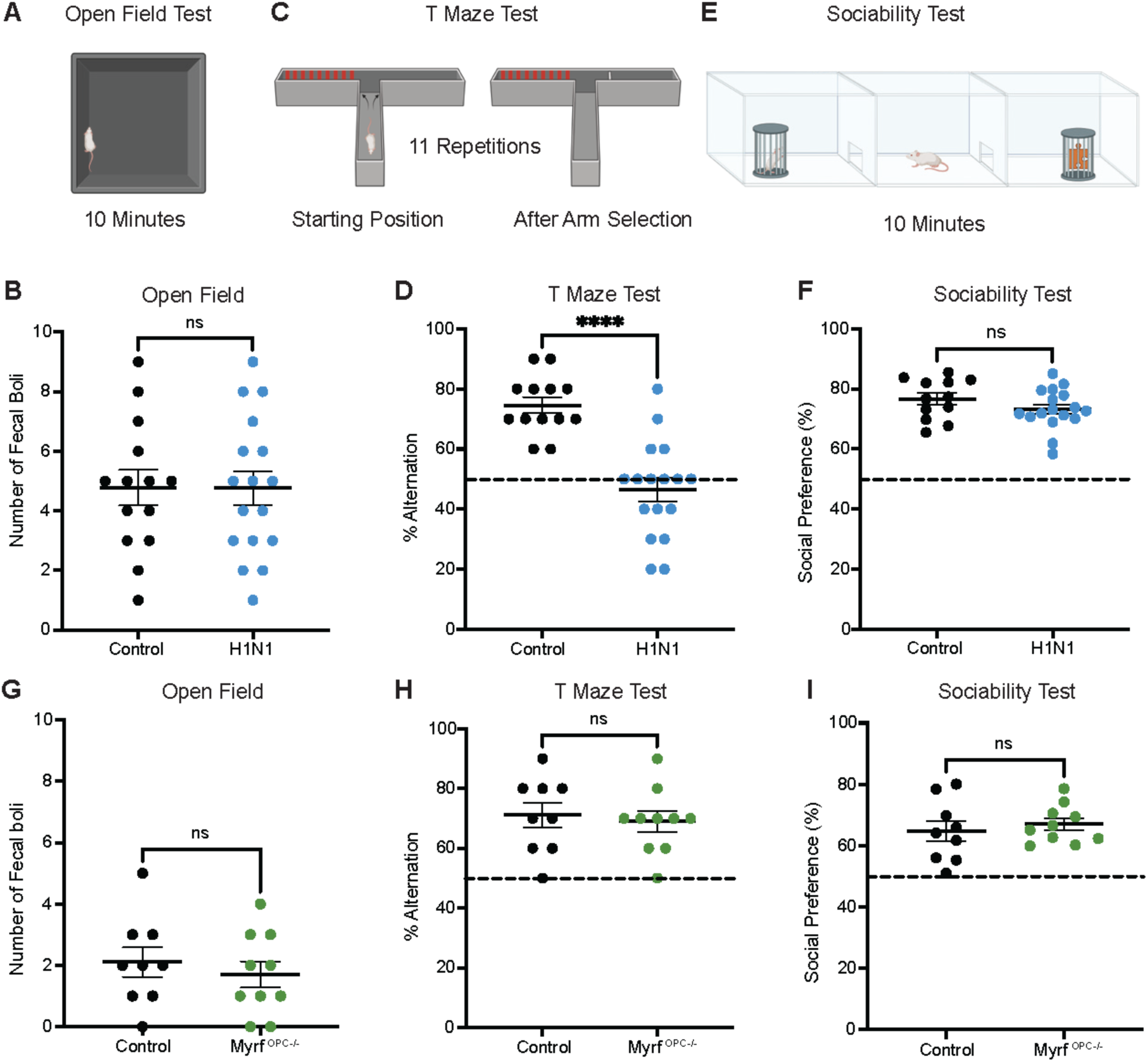
Hippocampal dependent cognitive deficits observed at four weeks post-infection, related to Figure 2. (A) Schematic illustration of the open field to assess anxiety-like behavior. Number of fecal boli in arena was quantified following a 10-minute testing period. (B) Schematic illustration of spontaneous T-maze alternation test to assess spatial working memory and hippocampal-related cognitive dysfunction. Percent alternation is calculated over 11 trials, allowing for 10 decisions. (C) Schematic illustration of three-chamber sociability test to assess social behavior and preference for social interaction. Social preference is defined as the percentage of time spent interacting with the conspecific mouse relative to total time spent interacting with either mouse or object. (D) Total number of fecal boli in the open field over 10-minute testing period. Open field performed at 4 weeks post-infection. *n* = 14 control, *n* = 17 H1N1 mice. (E) Percentage of alternations in T-maze performed at 4 weeks post-infection. *n* = 13 control, *n* = 17 H1N1 mice. (F) Sociability test performed at 4 weeks post-infection over 10-minute testing period. *n* = 12 control, *n* = 17 H1N1 mice. (G) Total number of fecal boli in the open field over 10-minute testing period. Open field performed at P42, 4 weeks post-injection. *n* = 7 control, *n* = 6 *Myrf^OPC-/-^* mice. (H) Percentage of alternations in T-maze performed at P42, 4 weeks post-injection. *n* = 7 control, *n* = 6 *Myrf^OPC-/-^* mice. (I) Sociability test performed at 4 weeks post-infection over 10-minute testing period. *n* = 7 control, *n* = 6 *Myrf^OPC-/-^* mice. Data shown as mean ± SEM; each dot represents an individual mouse. ns: p > 0.05, **** p < 0.0001, analyzed via (D-G, I) unpaired two-tailed t-test or (B and H) Mann-Whitney U test for datasets that failed the Shapiro-Wilk test.

**Figure S4.**
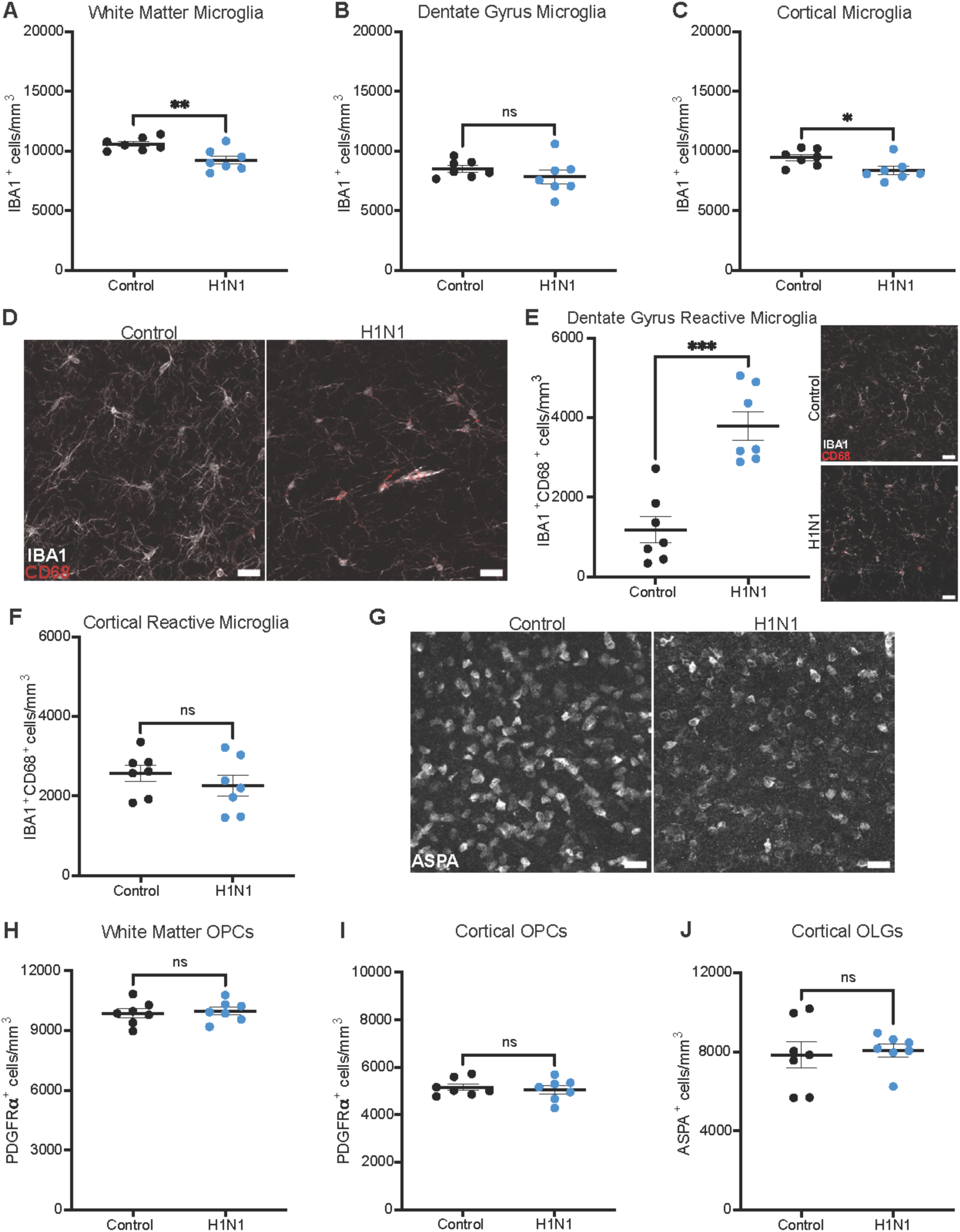
Glial cell counts 4 weeks post-infection in the white matter, dentate gyrus, and cortex, related to Figure 2. (A) Quantification of white matter microglia (IBA1^+^) 4 weeks post-infection in the cingulum. *n* = 7 mice per group. (B) Quantification of white matter microglia (IBA1^+^) 4 weeks post-infection in the hilus of the dentate gyrus. *n* = 7 mice per group. (C) Quantification of gray matter microglia (IBA1^+^) 4 weeks post-infection in the M2 region of cortex. *n* = 7 mice per group. (D) Representative confocal micrographs of white matter reactive microglia (IBA1, white; CD68, red) in the cingulum 4 weeks post-infection. (E) Quantification and representative confocal micrographs of white matter reactive microglia (IBA1^+^CD68^+^) 4 weeks post-infection in the hilus of the dentate gyrus. *n* = 7 mice per group. (F) Quantification of gray matter reactive microglia (IBA1^+^ CD68^+^) 4 weeks post-infection in the M2 region of cortex. *n* = 7 mice per group. (G) Representative confocal micrographs of white matter mature oligodendrocytes (ASPA, white) in the cingulum 4 weeks post-infection. (H) Quantification of white matter OPCs (PDGFRα^+^) 4 weeks post-infection in the cingulum. *n* = 7 mice per group. (I) Quantification of gray matter OPCs (PDGFRα^+^) 4 weeks post-infection in the M2 region of cortex. *n* = 7 mice per group. (J) Quantification of gray matter mature oligodendrocytes (ASPA^+^) 4 weeks post-infection in the M2 region of cortex. *n* = 7 mice per group. Data shown as mean ± SEM; each dot represents an individual mouse. ns: p > 0.05, * p < 0.05, ** p < 0.01, *** p < 0.001, analyzed via unpaired two-tailed t-test. (D, E, and G) Scale bars, 40 μm.

**Figure S5.**
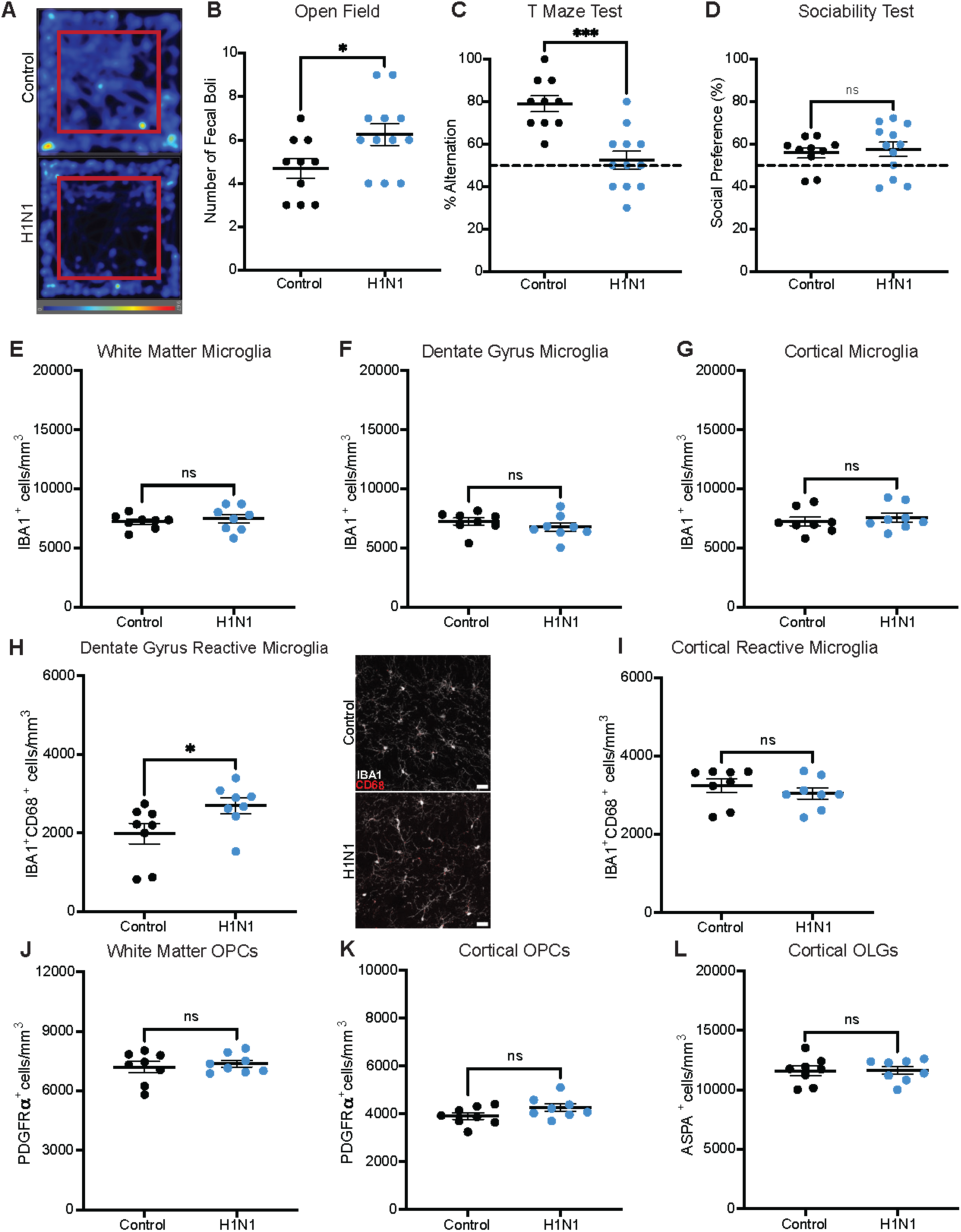
Impact of juvenile H1N1 infection on glia and behavior at 8 weeks, related to Figure 3. (A) Representative heatmaps illustrating time spent in the center (red square) and periphery of the arena during the open field test, with color intensity normalized across all trials (one mouse per trial). (B) Total number of fecal boli in the open field over 10-minute testing period. Open field performed at 8 weeks post-infection. *n* = 10 control, *n* = 12 H1N1 mice. (C) Percentage of alternations in T-maze performed at 8 weeks post-infection. *n* = 10 control, *n* = 12 H1N1 mice. (D) Sociability test performed over 10-minute testing period at 8 weeks post-infection. *n* = 10 control, *n* = 12 H1N1 mice. (E) Quantification of white matter microglia (IBA1^+^) 8 weeks post-infection in the cingulum. *n* = 8 mice per group. (F) Quantification of white matter microglia (IBA1^+^) 8 weeks post-infection in the hilus of the dentate gyrus. *n* = 8 mice per group. (G) Quantification of cortical gray matter microglia (IBA1^+^) 8 weeks post-infection in the M2 region of cortex. *n* = 8 mice per group. (H) Quantification and representative confocal micrographs of white matter reactive microglia (IBA1^+^CD68^+^) 8 weeks post-infection in the hilus of the dentate gyrus. *n* = 8 mice per group. (I) Quantification of cortical gray matter reactive microglia (IBA1^+^ CD68^+^) 8 weeks post-infection in the M2 region of cortex. *n* = 8 mice per group. (J) Quantification of white matter OPCs (PDGFRα^+^) 8 weeks post-infection in the cingulum. *n* = 8 mice per group. (K) Quantification of gray matter OPCs (PDGFRα^+^) 8 weeks post-infection in the M2 region of cortex. *n* = 8 mice per group. (L) Quantification of gray matter mature oligodendrocytes (ASPA^+^) 8 weeks post-infection in the M2 region of cortex. *n* = 8 mice per group. Data shown as mean ± SEM; each dot represents an individual mouse. ns: p > 0.05, * p < 0.05, *** p < 0.001, analyzed via (B, C, E-H, J-L) unpaired two-tailed t-test or (D and I) Mann-Whitney U test for datasets that failed the Shapiro-Wilk test. (G) Scale bars, 40 μm.

**Figure S6.**
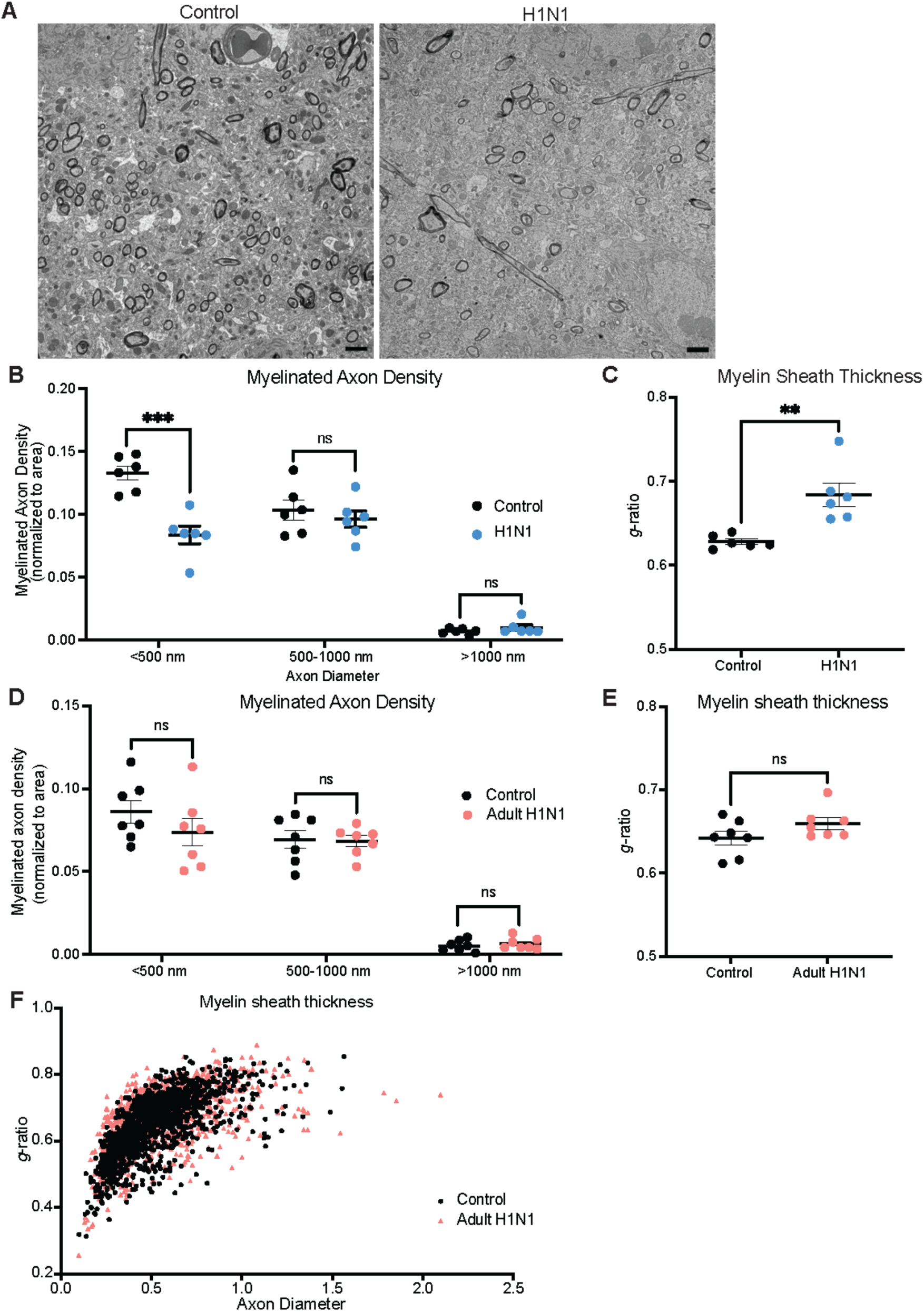
Juvenile H1N1 infection causes a decrease in myelinated axon density specifically in small diameter axons, related to Figure 3. (A) Representative TEM images at the level of the cingulum of the corpus callosum in cross-section in mice infected during the juvenile period. Myelinated axons visible as electron-dense sheaths encircling axons. (B) Myelinated axon density relative to small (<500 nm), medium (500-1000 nm), and large (>1000 nm) caliber axons for control (black points) or H1N1 (blue points) treated mice infected during the juvenile period 8 weeks post-infection. *n* = 6 mice per group. (C) Quantification of *g*-ratio of mice infected during the juvenile period 8 weeks post-infection. *n* = 6 mice per group. (D) Myelinated axon density relative to small (<500 nm), medium (500-1000 nm), and large (>1000 nm) caliber axons for control (black points) or H1N1 (orange points) treated mice injected during adulthood. *n* = 6 mice per group. (E) Quantification of *g*-ratio of mice infected during the juvenile period. *n* = 7 mice per group. (F) Scatter plots of *g*-ratio as a function of axon diameter 8 weeks post-infection for control axons (black dots) or H1N1 axons (orange triangles). Mice were infected during adulthood. A single point indicates the *g*-ratio for a single axon. Approximately 150-250 axons were quantified for each animal. *n* = 7 mice per group. Data shown as mean ± SEM; each dot represents an individual mouse. ns: p > 0.05, ** p < 0.01, *** p < 0.00, analyzed via (B) unpaired two-tailed t-test with correction for multiple comparisons or (C) unpaired two-tailed t-test. (A) Scale bars, 2 μm.

**Figure S7.**
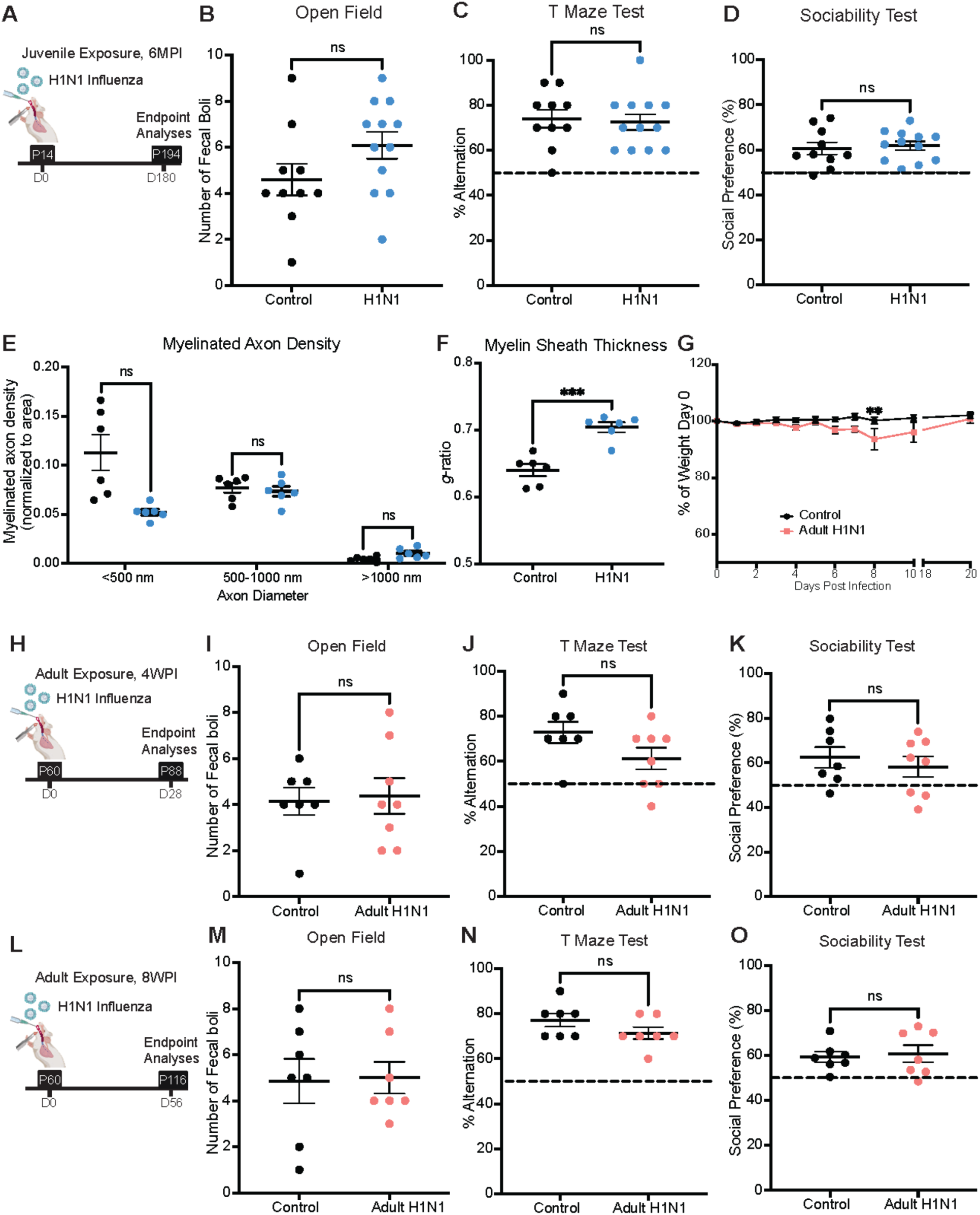
Behavioral assessments of anxiety-like behavior, sociability, and hippocampal function at chronic timepoints following juvenile and adult H1N1 infection, related to Figure 4. (A) Schematic illustration of experimental timeline. Mice were intratracheally inoculated at P14 with PBS or H1N, and endpoint analyses were performed at P194, 6 months post-infection (MPI). (B) Total number of fecal boli in the open field over 10-minute testing period. Open field performed at 6 months post-infection. *n* = 10 control, *n* = 12 H1N1 mice. (C) Percentage of alternations in T-maze performed at 6 months post-infection. *n* = 10 control, *n* = 12 H1N1 mice. (D) Sociability test performed over 10-minute testing period at 6 months post-infection. *n* = 10 control, *n* = 12 H1N1 mice. (E) Myelinated axon density relative to small (<500 nm), medium (500-1000 nm), and large (>1000 nm) caliber axons for control (black points) or H1N1 (blue points) treated mice infected during the juvenile period 6 months post-infection. *n* = 6 mice per group. (F) Quantification of *g*-ratio of mice infected during the juvenile period 6 months post-infection. *n* = 6 mice per group. (G) Body weight (% of day 0 weight) of control and H1N1 influenza mice infected during adulthood. *n* = 7-10 mice per group per timepoint. (H) Schematic illustration of experimental timeline. Mice were intratracheally inoculated at P60 with PBS or H1N1, and endpoint analyses were performed at P88, 4 weeks post-infection. (I) Total number of fecal boli in the open field. Open field performed at 4 weeks post-infection. *n* = 7 control, *n* = 8 H1N1 mice. (J) Percentage of alternations in T-maze performed at 4 weeks post-infection. *n* = 7 control, *n* = 8 H1N1 mice. (K) Sociability test performed at 4 weeks post-infection. *n* = 7 control, *n* = 8 H1N1 mice. (L) Schematic illustration of experimental timeline. Mice were intratracheally inoculated at P60 with PBS or H1N1, and endpoint analyses performed at P116, 8 weeks post-infection. (M) Total number of fecal boli in the open field. Open field performed at 8 weeks post-infection. *n* = 7 mice per group. (N) T-maze performed at 8 weeks post-infection. *n* = 7 mice per group. (O) Sociability test performed at 8 weeks post-infection. *n* = 7 mice per group. Data shown as mean ± SEM; each dot represents an individual mouse. ns: p > 0.05, analyzed via (B, D, F, I-K, M-O) unpaired two-tailed t-test, (C) Mann-Whitney U test for datasets that failed the Shapiro-Wilk test, or (E) unpaired two-tailed t-test with correction for multiple comparisons, or (G) two-way ANOVA with multiple comparisons.

**Figure S8.**
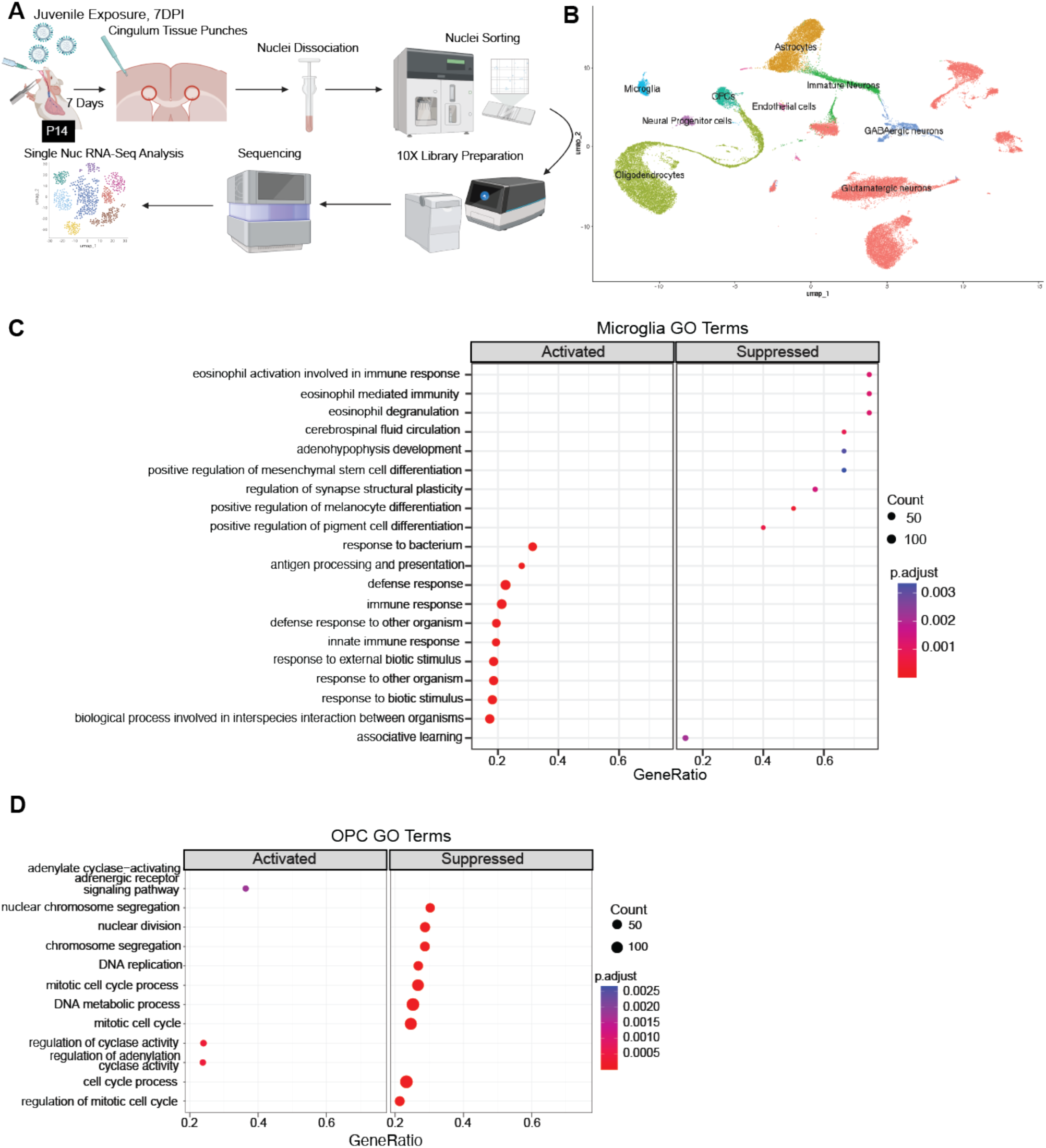
H1N1 alters transcriptional profiles of microglia and OPCs in the cingulum, related to Figure 5. (A) Schematic of experimental paradigm, sample collection and processing. Mice were intratracheally inoculated with PBS or H1N1 at P14, and brains were collected 7 days post-infection. Cingulum tissue punches were collected, nuclei were dissociated and sorted, and then libraries were prepared to perform single-nucleus RNA sequencing and analysis. *n* = 3 mice per group. (B) UMAP plot of 54,226 single nuclei from the cingulum, divided between control and H1N1 samples (26,653 single nuclei from control mice and 27,573 single nuclei from H1N1 mice). (C) Gene ontology (GO) enrichment analysis of differentially expressed genes in microglia cluster of H1N1 compared to control samples. Bubble plot displays the top enriched biological process (BP) GO terms for marker genes identified via single-nucleus RNA sequencing, The x-axis indicates the number of genes associated with the GO term divided by the total number of genes in the cluster. The bubble size is proportional to the number of genes enriched in each GO term, while the color represents the adjusted p-value. (D) GO enrichment analysis of differentially expressed genes in OPC cluster of H1N1 compared to control samples.

**Figure S9.**
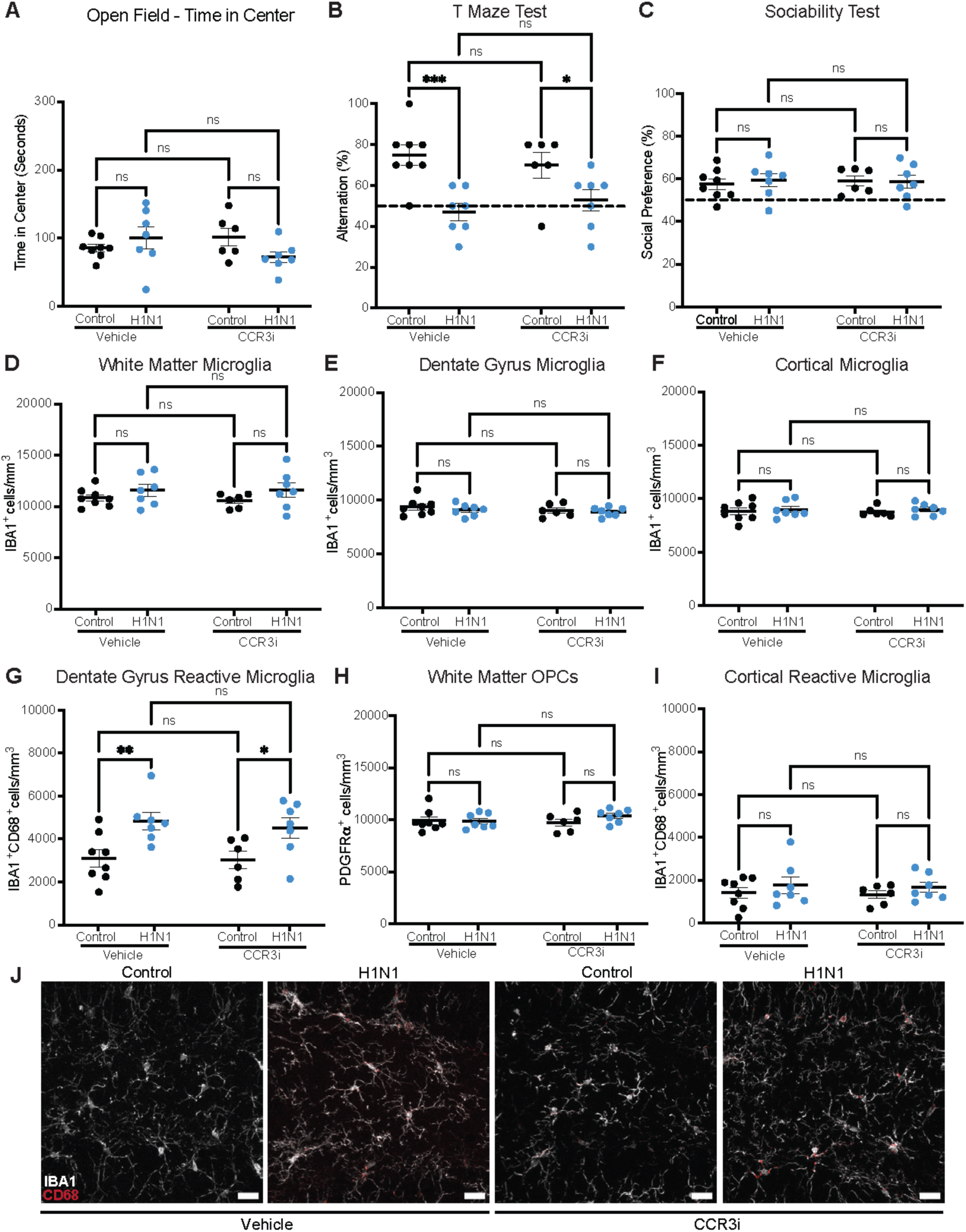
Chemokine receptor inhibition does not rescue hippocampal microglial reactivity or deficits in spontaneous alternating T-maze, related to Figure 6. (A) Time spent in center of the open field over 10-minute testing period. *n* = 8, 7, 6, and 7 mice per group. (B) Percentage of alternations in T-maze over 11 trials. *n* = 8, 7, 6, and 7 mice per group. (C) Results for sociability test performed over 10-minute testing period. *n* = 8, 7, 6, and 7 mice per group. (D) Quantification of white matter microglia (IBA1^+^) in the cingulum. *n* = 8, 7, 6, and 7 mice per group. (E) Quantification of white matter microglia (IBA1^+^) in the hilus of the dentate gyrus. *n* = 8, 7, 6, and 7 mice per group. (F) Quantification of gray matter microglia (IBA1^+^) in the M2 region of cortex. *n* = 8, 7, 6, and 7 mice per group. (G) Quantification of white matter OPCs (PDGFRα^+^) in the cingulum. *n* = 8, 7, 6, and 7 mice per group. (H) Quantification of white matter reactive microglia (IBA1^+^CD68^+^) in the hilus of the dentate gyrus. *n* = 8, 7, 6, and 7 mice per group. (I) Quantification of gray matter reactive microglia (IBA1^+^ CD68^+^) in the M2 region of cortex. *n* = 8, 7, 6, and 7 mice per group. (J) Representative confocal micrographs of reactive microglia (IBA1, white; CD68, red) in the hilus of the dentate gyrus. Data shown as mean ± SEM; each dot represents an individual mouse. ns: p > 0.05, * p < 0.05, ** p < 0.01, analyzed via two-way ANOVA. (J) Scale bars, 40 μm.

## REFERENCES

1. Gibson, E.M., Nagaraja, S., Ocampo, A., Tam, L.T., Wood, L.S., Pallegar, P.N., Greene, J.J., Geraghty, A.C., Goldstein, A.K., Ni, L., et al. (2019). Methotrexate Chemotherapy Induces Persistent Tri-glial Dysregulation that Underlies Chemotherapy-Related Cognitive Impairment. Cell 176, 43–55.e13. 10.1016/j.cell.2018.10.049.

2. Fernández-Castañeda, A., Lu, P., Geraghty, A.C., Song, E., Lee, M.-H., Wood, J., O’Dea, M.R., Dutton, S., Shamardani, K., Nwangwu, K., et al. (2022). Mild respiratory COVID can cause multi-lineage neural cell and myelin dysregulation. Cell, S0092867422007139. 10.1016/j.cell.2022.06.008.

3. Geraghty, A.C., Acosta-Alvarez, L., Rotiroti, M.C., Dutton, S., O’Dea, M.R., Kim, W., Trivedi, V., Mancusi, R., Shamardani, K., Malacon, K., et al. (2025). Immunotherapy-related cognitive impairment after CAR T cell therapy in mice. Cell 188, 3238–3258.e25. 10.1016/j.cell.2025.03.041.

4. Geraghty, A.C., Gibson, E.M., Ghanem, R.A., Greene, J.J., Ocampo, A., Goldstein, A.K., Ni, L., Yang, T., Marton, R.M., Paşca, S.P., et al. (2019). Loss of Adaptive Myelination Contributes to Methotrexate Chemotherapy-Related Cognitive Impairment. Neuron 103, 250–265.e8. 10.1016/j.neuron.2019.04.032.

5. Monje, M.L., Mizumatsu, S., Fike, J.R., and Palmer, T.D. (2002). Irradiation induces neural precursor-cell dysfunction. Nat. Med. 8, 955–962. 10.1038/nm749.

6. Monje, M.L., Toda, H., and Palmer, T.D. (2003). Inflammatory Blockade Restores Adult Hippocampal Neurogenesis. Science 302, 1760–1765. 10.1126/science.1088417.

7. Monje, M.L., Vogel, H., Masek, M., Ligon, K.L., Fisher, P.G., and Palmer, T.D. (2007). Impaired human hippocampal neurogenesis after treatment for central nervous system malignancies. Ann. Neurol. 62, 515–520. 10.1002/ana.21214.

8. Miron, V.E., Boyd, A., Zhao, J.-W., Yuen, T.J., Ruckh, J.M., Shadrach, J.L., Van Wijngaarden, P., Wagers, A.J., Williams, A., Franklin, R.J.M., et al. (2013). M2 microglia and macrophages drive oligodendrocyte differentiation during CNS remyelination. Nat. Neurosci. 16, 1211–1218. 10.1038/nn.3469.

9. McNamara, N.B., Munro, D.A.D., Bestard-Cuche, N., Uyeda, A., Bogie, J.F.J., Hoffmann, A., Holloway, R.K., Molina-Gonzalez, I., Askew, K.E., Mitchell, S., et al. (2023). Microglia regulate central nervous system myelin growth and integrity. Nature 613, 120–129. 10.1038/s41586-022-05534-y.

10. Liddelow, S.A., Guttenplan, K.A., Clarke, L.E., Bennett, F.C., Bohlen, C.J., Schirmer, L., Bennett, M.L., Münch, A.E., Chung, W.-S., Peterson, T.C., et al. (2017). Neurotoxic reactive astrocytes are induced by activated microglia. Nature 541, 481–487. 10.1038/nature21029.

11. Monje, M., and Iwasaki, A. (2022). The neurobiology of long COVID. Neuron 110, 3484–3496. 10.1016/j.neuron.2022.10.006.

12. VanRyzin, J.W., Pickett, L.A., and McCarthy, M.M. (2018). Microglia: Driving critical periods and sexual differentiation of the brain. Dev. Neurobiol. 78, 580–592. 10.1002/dneu.22569.

13. Hensch, T.K. (2004). Critical Period Regulation. Annu Rev Neurosci, 549–579. 10.1146/annurev.neuro.27.070203/144327.

14. Fletcher, J.L., Makowiecki, K., Cullen, C.L., and Young, K.M. (2021). Oligodendrogenesis and myelination regulate cortical development, plasticity and circuit function. Semin. Cell Dev. Biol. 118, 14–23. 10.1016/j.semcdb.2021.03.017.

15. Steadman, P.E., Xia, F., Ahmed, M., Mocle, A.J., Penning, A.R.A., Geraghty, A.C., Steenland, H.W., Monje, M., Josselyn, S.A., and Frankland, P.W. (2020). Disruption of Oligodendrogenesis Impairs Memory Consolidation in Adult Mice. Neuron 105, 150–164.e6. 10.1016/j.neuron.2019.10.013.

16. Allen, N.J., and Lyons, D.A. (2018). Glia as architects of central nervous system formation and function. Science 362, 181–185. 10.1126/science.aat0473.

17. Almeida, R.G., and Lyons, D.A. (2017). On Myelinated Axon Plasticity and Neuronal Circuit Formation and Function. J. Neurosci. 37, 10023–10034. 10.1523/JNEUROSCI.3185-16.2017.

18. Depp, C., Sun, T., Sasmita, A.O., Spieth, L., Berghoff, S.A., Nazarenko, T., Overhoff, K., Steixner-Kumar, A.A., Subramanian, S., Arinrad, S., et al. (2023). Myelin dysfunction drives amyloid-β deposition in models of Alzheimer’s disease. Nature 618, 349–357. 10.1038/s41586-023-06120-6.

19. Sasmita, A.O., Depp, C., Nazarenko, T., Sun, T., Siems, S.B., Ong, E.C., Nkeh, Y.B., Böhler, C., Yu, X., Bues, B., et al. (2024). Oligodendrocytes produce amyloid-β and contribute to plaque formation alongside neurons in Alzheimer’s disease model mice. Nat. Neurosci. 27, 1668–1674. 10.1038/s41593-024-01730-3.

20. Nave, K.-A., and Ehrenreich, H. (2014). Myelination and Oligodendrocyte Functions in Psychiatric Diseases. JAMA Psychiatry 71, 582. 10.1001/jamapsychiatry.2014.189.

21. Yakovlev, P., and Lecours, A.-R. (1967). The myelogenetic cycles of regional maturation of the brain. In Regional development of the brain in early life (Blackwell Scientific), pp. 3–70.

22. Lebel, C., Gee, M., Camicioli, R., Wieler, M., Martin, W., and Beaulieu, C. (2012). Diffusion tensor imaging of white matter tract evolution over the lifespan. NeuroImage 60, 340–352. 10.1016/j.neuroimage.2011.11.094.

23. Semple, B.D., Blomgren, K., Gimlin, K., Ferriero, D.M., and Noble-Haeusslein, L.J. (2013). Brain development in rodents and humans: Identifying benchmarks of maturation and vulnerability to injury across species. Prog. Neurobiol. 106–107, 1–16. 10.1016/j.pneurobio.2013.04.001.

24. Cristobal, C.D., and Lee, H.K. (2022). Development of myelinating glia: An overview. Glia 70, 2237–2259. 10.1002/glia.24238.

25. Baum, G.L., Flournoy, J.C., Glasser, M.F., Harms, M.P., Mair, P., Sanders, A.F.P., Barch, D.M., Buckner, R.L., Bookheimer, S., Dapretto, M., et al. (2022). Graded Variation in T1w/T2w Ratio during Adolescence: Measurement, Caveats, and Implications for Development of Cortical Myelin. J. Neurosci. 42, 5681–5694. 10.1523/JNEUROSCI.2380-21.2022.

26. Murakami, J.W., Weinberger, E., and Shaw, D.W.W. Normal Myelination of the Pediatric Brain Imaged with Fluid-Attenuated Inversion-Recovery (FLAIR) MR Imaging.

27. Dietrich, R., Bradley, W., Zaragoza, E., Otto, R., Taira, R., Wilson, G., and Kangarloo, H. (1988). MR evaluation of early myelination patterns in normal and developmentally delayed infants. Am. J. Roentgenol. 150, 889–896. 10.2214/ajr.150.4.889.

28. Hildebrand, C., Remahl, S., Persson, H., and Bjartmar, C. (1993). Myelinated nerve fibres in the CNS. Prog. Neurobiol. 40, 319–384. 10.1016/0301-0082(93)90015-K.

29. Almeida, R.G. (2018). The Rules of Attraction in Central Nervous System Myelination. Front. Cell. Neurosci. 12, 367. 10.3389/fncel.2018.00367.

30. Shi, L., Fatemi, S.H., Sidwell, R.W., and Patterson, P.H. (2003). Maternal Influenza Infection Causes Marked Behavioral and Pharmacological Changes in the Offspring. J. Neurosci. 23, 297–302. 10.1523/JNEUROSCI.23-01-00297.2003.

31. Fatemi, S.H., Pearce, D.A., Brooks, A.I., and Sidwell, R.W. (2005). Prenatal viral infection in mouse causes differential expression of genes in brains of mouse progeny: A potential animal model for schizophrenia and autism. Synapse 57, 91–99. 10.1002/syn.20162.

32. Haddad, F.L., Patel, S.V., and Schmid, S. (2020). Maternal Immune Activation by Poly I:C as a preclinical Model for Neurodevelopmental Disorders: A focus on Autism and Schizophrenia. Neurosci. Biobehav. Rev. 113, 546–567. 10.1016/j.neubiorev.2020.04.012.

33. Smith, A.P. (2013). Twenty-five years of research on the behavioural malaise associated with influenza and the common cold. Psychoneuroendocrinology 38, 744–751. 10.1016/j.psyneuen.2012.09.002.

34. Morales-Nebreda, L., Chi, M., Lecuona, E., Chandel, N.S., Dada, L.A., Ridge, K., Soberanes, S., Nigdelioglu, R., Sznajder, J.I., Mutlu, G.M., et al. (2014). Intratracheal administration of influenza virus is superior to intranasal administration as a model of acute lung injury. J. Virol. Methods 209, 116–120. 10.1016/j.jviromet.2014.09.004.

35. Song, E., Zhang, C., Israelow, B., Lu-Culligan, A., Prado, A.V., Skriabine, S., Lu, P., Weizman, O.-E., Liu, F., Dai, Y., et al. (2021). Neuroinvasion of SARS-CoV-2 in human and mouse brain. J. Exp. Med. 218, e20202135. 10.1084/jem.20202135.

36. Jurgens, H.A., Amancherla, K., and Johnson, R.W. (2012). Influenza Infection Induces Neuroinflammation, Alters Hippocampal Neuron Morphology, and Impairs Cognition in Adult Mice. J. Neurosci. 32, 3958–3968. 10.1523/JNEUROSCI.6389-11.2012.

37. Jurgens, H.A., and Johnson, R.W. (2012). Environmental enrichment attenuates hippocampal neuroinflammation and improves cognitive function during influenza infection. Brain. Behav. Immun. 26, 1006–1016. 10.1016/j.bbi.2012.05.015.

38. Bronner, M.B., Knoester, H., Sol, J.J., Bos, A.P., Heymans, H.S.A., and Grootenhuis, M.A. (2009). An explorative study on quality of life and psychological and cognitive function in pediatric survivors of septic shock: Pediatr. Crit. Care Med. 10, 636–642. 10.1097/PCC.0b013e3181ae5c1a.

39. Kaur, J., Singhi, P., Singhi, S., Malhi, P., and Saini, A.G. (2016). Neurodevelopmental and Behavioral Outcomes in Children With Sepsis-Associated Encephalopathy Admitted to Pediatric Intensive Care Unit: A Prospective Case Control Study. J. Child Neurol. 31, 683–690. 10.1177/0883073815610431.

40. Crivelli, L., Calandri, I., Corvalán, N., Carello, M.A., Keller, G., Martínez, C., Arruabarrena, M., and Allegri, R. (2022). Cognitive consequences of COVID-19: results of a cohort study from South America. Arq. Neuropsiquiatr. 80, 240–247. 10.1590/0004-282x-anp-2021-0320.

41. Panagea, E., Messinis, L., Petri, M.C., Liampas, I., Anyfantis, E., Nasios, G., Patrikelis, P., and Kosmidis, M. (2025). Neurocognitive Impairment in Long COVID: A Systematic Review. Arch. Clin. Neuropsychol. 40, 125–149. 10.1093/arclin/acae042.

42. Calsavara, A.J.C., Nobre, V., Barichello, T., and Teixeira, A.L. (2018). Post-sepsis cognitive impairment and associated risk factors: A systematic review. Aust. Crit. Care 31, 242–253. 10.1016/j.aucc.2017.06.001.

43. Leger, M., Quiedeville, A., Bouet, V., Haelewyn, B., Boulouard, M., Schumann-Bard, P., and Freret, T. (2013). Object recognition test in mice. Nat. Protoc. 8, 2531–2537. 10.1038/nprot.2013.155.

44. Antunes, M., and Biala, G. (2012). The novel object recognition memory: neurobiology, test procedure, and its modifications. Cogn. Process. 13, 93–110. 10.1007/s10339-011-0430-z.

45. Deacon, R.M.J., and Rawlins, J.N.P. (2006). T-maze alternation in the rodent. Nat. Protoc. 1, 7–12. 10.1038/nprot.2006.2.

46. d’Isa, R., Comi, G., and Leocani, L. (2021). Apparatus design and behavioural testing protocol for the evaluation of spatial working memory in mice through the spontaneous alternation T-maze. Sci. Rep. 11, 21177. 10.1038/s41598-021-00402-7.

47. McKenzie, I.A., Ohayon, D., Li, H., Paes De Faria, J., Emery, B., Tohyama, K., and Richardson, W.D. (2014). Motor skill learning requires active central myelination. Science 346, 318–322. 10.1126/science.1254960.

48. Rao, S., Gross, R.S., Mohandas, S., Stein, C.R., Case, A., Dreyer, B., Pajor, N.M., Bunnell, H.T., Warburton, D., Berg, E., et al. (2024). Postacute Sequelae of SARS-CoV-2 in Children. Pediatrics 153, e2023062570. 10.1542/peds.2023-062570.

49. Foret-Bruno, P., Shafran, R., Stephenson, T., Nugawela, M.D., Chan, D., Ladhani, S., McOwat, K., Mensah, A., Simmons, R., Fox Smith, L., et al. (2024). Prevalence and co-occurrence of cognitive impairment in children and young people up to 12-months post infection with SARS-CoV-2 (Omicron variant). Brain. Behav. Immun. 119, 989–994. 10.1016/j.bbi.2024.05.001.

50. Gross, R.S., Thaweethai, T., Kleinman, L.C., Snowden, J.N., Rosenzweig, E.B., Milner, J.D., Tantisira, K.G., Rhee, K.E., Jernigan, T.L., Kinser, P.A., et al. (2024). Characterizing Long COVID in Children and Adolescents. JAMA 332, 1174. 10.1001/jama.2024.12747.

51. Keren-Shaul, H., Spinrad, A., Weiner, A., Matcovitch-Natan, O., Dvir-Szternfeld, R., Ulland, T.K., David, E., Baruch, K., Lara-Astaiso, D., Toth, B., et al. (2017). A Unique Microglia Type Associated with Restricting Development of Alzheimer’s Disease. Cell 169, 1276–1290.e17. 10.1016/j.cell.2017.05.018.

52. Silvin, A., Uderhardt, S., Piot, C., Da Mesquita, S., Yang, K., Geirsdottir, L., Mulder, K., Eyal, D., Liu, Z., Bridlance, C., et al. (2022). Dual ontogeny of disease-associated microglia and disease inflammatory macrophages in aging and neurodegeneration. Immunity 55, 1448–1465.e6. 10.1016/j.immuni.2022.07.004.

53. Marques, S., Zeisel, A., Codeluppi, S., Van Bruggen, D., Mendanha Falcão, A., Xiao, L., Li, H., Häring, M., Hochgerner, H., Romanov, R.A., et al. (2016). Oligodendrocyte heterogeneity in the mouse juvenile and adult central nervous system. Science 352, 1326–1329. 10.1126/science.aaf6463.

54. Kenigsbuch, M., Bost, P., Halevi, S., Chang, Y., Chen, S., Ma, Q., Hajbi, R., Schwikowski, B., Bodenmiller, B., Fu, H., et al. (2022). A shared disease-associated oligodendrocyte signature among multiple CNS pathologies. Nat. Neurosci. 25, 876–886. 10.1038/s41593-022-01104-7.

55. Studahl, M. (2003). Influenza virus and CNS manifestations. J. Clin. Virol. 28, 225–232. 10.1016/S1386-6532(03)00119-7.

56. Surana, P., Tang, S., McDougall, M., Tong, C.Y.W., Menson, E., and Lim, M. (2011). Neurological complications of pandemic influenza A H1N1 2009 infection: European case series and review. Eur. J. Pediatr. 170, 1007–1015. 10.1007/s00431-010-1392-3.

57. Gross, R.S., Thaweethai, T., Salisbury, A.L., Kleinman, L.C., Mohandas, S., Rhee, K.E., Snowden, J.N., Tantisira, K.G., Warburton, D., Wood, J.C., et al. (2025). Characterizing Long COVID Symptoms During Early Childhood. JAMA Pediatr. 179, 781. 10.1001/jamapediatrics.2025.1066.

58. Toepfner, N., Brinkmann, F., Augustin, S., Stojanov, S., and Behrends, U. (2024). Long COVID in pediatrics—epidemiology, diagnosis, and management. Eur. J. Pediatr. 183, 1543–1553. 10.1007/s00431-023-05360-y.

59. Nayak, J., Hoy, G., and Gordon, A. (2021). Influenza in Children. Cold Spring Harb. Perspect. Med. 11, a038430. 10.1101/cshperspect.a038430.

60. CDC (2024). Disease Burden of Flu. https://www.cdc.gov/flu/about/burden/index.html.

61. CDC (2024). Flu & Children at Higher Risk. https://www.cdc.gov/flu/highrisk/children-high-risk.htm.

62. Garcia, A. (2022). “Tripledemic” of flu, RSV & COVID-19 cases continue to rise with Andrea Garcia, JD, MPH. Am. Med. Assoc. https://www.ama-assn.org/delivering-care/public-health/tripledemic-flu-rsv-covid-19-cases-continue-rise-andrea-garcia-jd-mph.

63. CDC Health Alert Network (2022). Increased Respiratory Virus Activity, Especially Among Children, Early in the 2022-2023 Fall and Winter (Centers for Disease Control and Prevention).

64. Riepl, A., Straßmayr, L., Voitl, P., Ehlmaier, P., Voitl, J.J.M., Langer, K., Kuzio, U., Mühl-Riegler, A., Mühl, B., and Diesner-Treiber, S.C. (2023). The surge of RSV and other respiratory viruses among children during the second COVID-19 pandemic winter season. Front. Pediatr. 11, 1112150. 10.3389/fped.2023.1112150.

65. Metz, C., Schmid, A., and Veldhoen, S. (2023). Increase in complicated upper respiratory tract infection in children during the 2022/2023 winter season—a post coronavirus disease 2019 effect? Pediatr. Radiol. 54, 49–57. 10.1007/s00247-023-05808-1.

66. Chuang, Y.-C., Lin, K.-P., Wang, L.-A., Yeh, T.-K., and Liu, P.-Y. (2023). The Impact of the COVID-19 Pandemic on Respiratory Syncytial Virus Infection: A Narrative Review. Infect. Drug Resist. Volume 16, 661–675. 10.2147/IDR.S396434.

67. Ravenholt, R.T., and Foege, W.H. (1982). 1918 Influenza, Encephalitis Lethargica, Parkinsonism. The Lancet 2, 860–864. 10.1016/s0140-6736(82)90820-0.%20PMID:%206126720.

68. Honigsbaum, M. (2013). “An inexpressible dread”: psychoses of influenza at fin-de-siècle. The Lancet 381, 988–989. 10.1016/S0140-6736(13)60701-1.

69. Menninger, K.A. (1919). Psychoses associated with influenza. JAMA 72, 235–241.

70. Auro, K., Holopainen, I., Perola, M., Havulinna, A.S., and Raevuori, A. (2024). Attention-Deficit/Hyperactivity Disorder Diagnoses in Finland During the COVID-19 Pandemic. JAMA Netw. Open 7, e2418204. 10.1001/jamanetworkopen.2024.18204.

71. Song, J., Park, S.J., Jeong, S., Chun, A.Y., and Park, S.M. (2025). Increasing incidence of ADHD among children, adolescents and young adults: COVID-19 pandemic-driven trend in Korea (2012–2023). BMJ Ment. Health 28, e301662. 10.1136/bmjment-2025-301662.

72. Sammons, M., Popescu, M.C., Chi, J., Liberles, S.D., Gogolla, N., and Rolls, A. (2024). Brain-body physiology: Local, reflex, and central communication. Cell 187, 5877–5890. 10.1016/j.cell.2024.08.050.

73. Waxman, S.G. (1980). Determinants of conduction velocity in myelinated nerve fibers. Muscle Nerve 3, 141–150. 10.1002/mus.880030207.

74. Fünfschilling, U., Supplie, L.M., Mahad, D., Boretius, S., Saab, A.S., Edgar, J., Brinkmann, B.G., Kassmann, C.M., Tzvetanova, I.D., Möbius, W., et al. (2012). Glycolytic oligodendrocytes maintain myelin and long-term axonal integrity. Nature 485, 517–521. 10.1038/nature11007.

75. Noori, R., Park, D., Griffiths, J.D., Bells, S., Frankland, P.W., Mabbott, D., and Lefebvre, J. (2020). Activity-dependent myelination: A glial mechanism of oscillatory self-organization in large-scale brain networks. Proc. Natl. Acad. Sci. 117, 13227–13237. 10.1073/pnas.1916646117.

76. Pajevic, S., Basser, P.J., and Fields, R.D. (2014). Role of myelin plasticity in oscillations and synchrony of neuronal activity. Neuroscience 276, 135–147. 10.1016/j.neuroscience.2013.11.007.

77. Gadani, S.P., Walsh, J.T., Smirnov, I., Zheng, J., and Kipnis, J. (2015). The Glia-Derived Alarmin IL-33 Orchestrates the Immune Response and Promotes Recovery following CNS Injury. Neuron 85, 703–709. 10.1016/j.neuron.2015.01.013.

78. Alves De Lima, K., Rustenhoven, J., Da Mesquita, S., Wall, M., Salvador, A.F., Smirnov, I., Martelossi Cebinelli, G., Mamuladze, T., Baker, W., Papadopoulos, Z., et al. (2020). Meningeal γδ T cells regulate anxiety-like behavior via IL-17a signaling in neurons. Nat. Immunol. 21, 1421–1429. 10.1038/s41590-020-0776-4.

79. Remsik, J., Wilcox, J.A., Babady, N.E., McMillen, T.A., Vachha, B.A., Halpern, N.A., Dhawan, V., Rosenblum, M., Iacobuzio-Donahue, C.A., Avila, E.K., et al. (2021). Inflammatory Leptomeningeal Cytokines Mediate COVID-19 Neurologic Symptoms in Cancer Patients. Cancer Cell 39, 276–283.e3. 10.1016/j.ccell.2021.01.007.

80. Acharya, M.M., Green, K.N., Allen, B.D., Najafi, A.R., Syage, A., Minasyan, H., Le, M.T., Kawashita, T., Giedzinski, E., Parihar, V.K., et al. (2016). Elimination of microglia improves cognitive function following cranial irradiation. Sci. Rep. 6, 31545. 10.1038/srep31545.

81. Andoh, M., and Koyama, R. (2021). Microglia regulate synaptic development and plasticity. Dev. Neurobiol. 81, 568–590. 10.1002/dneu.22814.

82. Paolicelli, R.C., Bolasco, G., Pagani, F., Maggi, L., Scianni, M., Panzanelli, P., Giustetto, M., Ferreira, T.A., Guiducci, E., Dumas, L., et al. (2011). Synaptic Pruning by Microglia Is Necessary for Normal Brain Development. Science 333, 1456–1458. 10.1126/science.1202529.

83. Schafer, D.P., Lehrman, E.K., Kautzman, A.G., Koyama, R., Mardinly, A.R., Yamasaki, R., Ransohoff, R.M., Greenberg, M.E., Barres, B.A., and Stevens, B. (2012). Microglia Sculpt Postnatal Neural Circuits in an Activity and Complement-Dependent Manner. Neuron 74, 691–705. 10.1016/j.neuron.2012.03.026.

84. Kopec, A.M., Smith, C.J., Ayre, N.R., Sweat, S.C., and Bilbo, S.D. (2018). Microglial dopamine receptor elimination defines sex-specific nucleus accumbens development and social behavior in adolescent rats. Nat. Commun. 9, 3769. 10.1038/s41467-018-06118-z.

85. Filipello, F., Morini, R., Corradini, I., Zerbi, V., Canzi, A., Michalski, B., Erreni, M., Markicevic, M., Starvaggi-Cucuzza, C., Otero, K., et al. (2018). The Microglial Innate Immune Receptor TREM2 Is Required for Synapse Elimination and Normal Brain Connectivity. Immunity 48, 979–991.e8. 10.1016/j.immuni.2018.04.016.

86. Schafer, D.P., Heller, C.T., Gunner, G., Heller, M., Gordon, C., Hammond, T., Wolf, Y., Jung, S., and Stevens, B. (2016). Microglia contribute to circuit defects in Mecp2 null mice independent of microglia-specific loss of Mecp2 expression. eLife 5, e15224. 10.7554/eLife.15224.

87. Kim, H.-J., Cho, M.-H., Shim, W.H., Kim, J.K., Jeon, E.-Y., Kim, D.-H., and Yoon, S.-Y. (2017). Deficient autophagy in microglia impairs synaptic pruning and causes social behavioral defects. Mol. Psychiatry 22, 1576–1584. 10.1038/mp.2016.103.

88. Block, C.L., Eroglu, O., Mague, S.D., Smith, C.J., Ceasrine, A.M., Sriworarat, C., Blount, C., Beben, K.A., Malacon, K.E., Ndubuizu, N., et al. (2022). Prenatal environmental stressors impair postnatal microglia function and adult behavior in males. Cell Rep. 40, 111161. 10.1016/j.celrep.2022.111161.

89. Guttenplan, K.A., Weigel, M.K., Prakash, P., Wijewardhane, P.R., Hasel, P., Rufen-Blanchette, U., Münch, A.E., Blum, J.A., Fine, J., Neal, M.C., et al. (2021). Neurotoxic reactive astrocytes induce cell death via saturated lipids. Nature 599, 102–107. 10.1038/s41586-021-03960-y.

90. Revelli, D.A., Boylan, J.A., and Gherardini, F.C. (2012). A non-invasive intratracheal inoculation method for the study of pulmonary melioidosis. Front. Cell. Infect. Microbiol. 2. 10.3389/fcimb.2012.00164.

91. Ianevski, A., Giri, A.K., and Aittokallio, T. (2022). Fully-automated and ultra-fast cell-type identification using specific marker combinations from single-cell transcriptomic data. Nat. Commun. 13, 1246. 10.1038/s41467-022-28803-w.

